# Evolution of ssDNA plant viruses in the natural environment - a journey through time

**DOI:** 10.1101/2025.09.10.675476

**Authors:** João P. Herrera da Silva, César A.D. Xavier, Phedra G.S. Oliveira, Márcio T. Godinho, Júlia B. Lage, José C.F. da Silva, Alison T.M. Lima, F. Murilo Zerbini

**Author notes:** Dep. of Veterinary Population Medicine, University of Minnesota, St. Paul, MN 55108, USA. Dep. of Entomology and Plant Pathology, North Carolina State University, Raleigh, NC, 27695, USA. Dep. de Microbiologia/BIOAGRO, Universidade Federal de Viçosa, Viçosa, MG 36570-900, Brazil.

## Abstract

Begomoviruses pose a major threat to food security, particularly in developing countries. These small ssDNA viruses exhibit substitution rates comparable to those of RNA viruses. The temporal dynamics of begomoviruses in non-agricultural environments have been largely overlooked, and little is known about how these viruses evolve in the absence of anthropogenic influence. In this study, we investigated the temporal dynamics of begomoviruses in a small fragment of regenerating Atlantic Forest area in the state of Minas Gerais, Brazil. Samples of *Sida acuta*, a wild plant native from South America, were collected at the same location over a 12-year period (2011-2022). Five distinct begomoviruses were detected infecting this host: OxYVV, SiYLCV, SimMV, MaYVV, and SiMV. OxYVV and SimMV were subdivided into multiple variants, revealing their potential as reservoirs of viral biodiversity. Several shifts in species and variant composition were observed in the viral community over time, with the most drastic change occurring in 2016, when SiYLCV outnumbered OxYVV. The reasons behind this turnover remain uncertain, but the most compelling clues point to a population expansion of SiYLCV. We detected a strong temporal signal in two of the most abundant viruses (OxYVV and SiYLCV), which allowed us to calibrate molecular clocks and estimate substitution rates for both of them. Our results indicate that, even in the natural environment, begomoviruses can evolve at rates similar to those reported in agricultural systems.

## Introduction

Begomoviruses (family *Geminiviridae*) are plant-infecting viruses with small, single-stranded, circular DNA genomes which can be mono- or bipartite (Fiallo-Olive et al., 2021). Monopartite begomoviruses, which are predominant in Europe, Africa, Asia and Oceania (EAAO), have genomes around 2.8 kilobases (kb) in length, while the bipartite begomoviruses, which are predominant in the Americas (AM), have two components, named DNA-A and DNA-B, each approximately 2.6 kb in length. The DNA-A and DNA-B differ from each other both in terms of gene content and in the function of those genes. The DNA-A encodes genes that are associated with viral replication, suppression of host defense responses, cell cycle reprogramming and particle assembly, while the DNA-B encodes genes which are associated with cell-to-cell and systemic movement in the plant. In the case of bipartite begomoviruses, both components are essential to trigger systemic infection (Hanley-Bowdoin et al., 2013). Begomoviruses have twinned, quasi-icosahedral particles formed by 110 subunits of a single structural (coat) protein. In the case of bipartite viruses, each component is encapsidated in a separate particle (Hesketh et al., 2018). They are transmitted in a persistent circulative manner by whiteflies of the *Bemisia tabaci* cryptic species complex (Campbell et al., 2023; De Barro et al., 2011).

Several epidemics caused by begomoviruses have been documented around the world, among them the epidemics of cassava mosaic in Sub-Saharan Africa, of tomato yellow leaf curl in the Mediterranean, of cotton leaf curl in the Indian Subcontinent, and of golden/yellow mosaic in legume, solanaceous and cucurbit crops in the Americas (Rojas et al., 2018). In all of these cases, a complex of closely related begomoviruses, including one or more recombinant viruses, were involved. The great diversity of begomoviruses is a result of their rapid evolutionary dynamics, driven by mutation rates which are equivalent to those of RNA viruses, as well as frequent recombination events and genomic reassortment (Duffy and Holmes, 2008; Duffy et al., 2008; Lima et al., 2017; Xavier et al., 2021).

Besides evolutionary mechanisms, ecological changes such as environmental disasters, ecosystem degradation and global connectivity can also affect the emergence of viruses. These events can contribute to the displacement of pathogens and/or their respective hosts and vectors to new niches, consequently increasing the chances of contact between viruses in non-cultivated and cultivated hosts and enabling the occurrence of spillover events that lead to the expansion of the virus host range (García-Arenal and Zerbini, 2019; Kilpatrick and Randolph, 2012; Pybus et al., 2015; Roossinck and Garcia-Arenal, 2015).

In Brazil, the emergence of begomoviruses in tomatoes during the 1990’s was driven by the introduction of *Bemisia tabaci* MEAM1 (then known as *B. tabaci* biotype B) into the country (Ribeiro et al., 1998). This polyphagous “supervector” facilitated the spillover of a large number of begomoviruses from their indigenous natural hosts to tomatoes, with at least two, tomato severe rugose virus (ToSRV) and tomato mottle leaf curl virus (ToMoLCV), becoming established as serious pathogens in that crop (Fernandes et al., 2008; Rocha et al., 2013; Souza et al., 2022).

A similar scenario was observed in the begomovirus epidemics in cassava in Africa during the 1980s and 1990s. Cassava is a plant native to South America, and there are no reports of begomoviruses infecting this crop in its region of origin. This suggests that the begomoviruses responsible for these epidemics were likely transmitted laterally from native African plants to cassava (Patil and Fauquet, 2009). Furthermore, the emergence of begomoviruses in African cassava was driven by viral synergistic interactions, recombination, and reassortments, which enhanced viral fitness in its new host (Pita et al., 2001). Similarly, several begomoviruses from the tomato yellow leaf curl complex emerged and expanded their host range through recombination, which facilitated their adaptation and spread to new host species (Belabess et al., 2016; Fiallo-Olivé et al., 2019).

The geographical range of viral hosts as well as a host’s permissiveness to infection by multiple viruses impacts the epidemiology of viral diseases (García-Arenal and Zerbini, 2019; Roossinck and Garcia-Arenal, 2015). In this sense, non-cultivated plants are essential players in the emergence of begomoviruses in agroecosystems. Some wild plants, if not managed, can remain in the field for a long period of time acting as a reservoir of viral diversity. Wild plants experience infection events by multiple viruses, and therefore constitute a favorable site for the occurrence of recombination events (Ferro et al., 2017a; García-Arenal and Zerbini, 2019; Lima et al., 2017; Rocha et al., 2013). Studies on begomovirus diversity in non-cultivated hosts have gained special attention in the 21st century (Castillo-Urquiza et al., 2008; Ferro et al., 2017b; Mar et al., 2017; Rodríguez-Negrete et al., 2019; Shakir et al., 2019). Most of these investigations have been conducted in the context of agricultural production. Understanding the evolutionary dynamics of plant viruses in natural (non-agricultural) ecosystems could provide insights into how these organisms evolve without the pressures imposed by anthropogenic activity, making it possible to draw parallels between these two contexts and eventually leading to more effective strategies to mitigate virus emergence in crops (Dolan et al., 2018; García-Arenal and Zerbini, 2019).

The spatio-temporal evolutionary dynamics of geminiviruses has been addressed in a number of studies and has expanded our understanding of the processes that contribute to the differentiation of these populations over the time, as well as estimating their evolutionary rates (Duffy and Holmes, 2009; Sánchez-Campos et al., 2002; Yang et al., 2014). However, as mentioned above, all these studies were carried out under an agricultural context. Moreover, most studies that explored temporal aspects spanned large areas, with some specific locations and/or time points being overrepresented in relation to others (Duffy and Holmes, 2009; Mar et al., 2017; Rodelo-Urrego et al., 2013).

In this work we explored the temporal dynamics of the begomovirus community infecting the non-cultivated plant *Sida acuta* (Malvaceae) in the natural environment (outside of an agricultural context). Samples were collected annually over 12 years at the same place and at the same time of year (late spring). We seek to understand how the diversity and variability of begomoviruses are distributed in a small host population and whether they reflect the patterns observed in larger areas. Furthermore, we explored aspects associated with temporal diversification, seeking to understand how these viruses evolve under low anthropogenic influence compared to agroecosystems.

## Materials and Methods

### Data sampling

*Sida acuta* plants (Malvaceae) displaying symptoms of begomovirus infection (Suppl. Figure S1) were collected annually in early December over a 12-year period (2011-2022). The sampling area, approximately 300 m², is located in a transition zone between Atlantic Forest remnants and a suburban area outside of an agricultural context, with minimal human intervention, near Viçosa, Minas Gerais, Brazil (−20.780960, −42.883434; Suppl. Figure S2). Whiteflies were sporadically present in the area, in much lower numbers compared to what is normally observed in agricultural fields. On average, 35 plants were collected per year, totaling 360 plants over the study period. Each specimen was identified, photographed, press-dried, and stored at room temperature in the Laboratory of Virus Ecology and Evolution at UFV.

### Cloning and sequencing

Total DNA was extracted from plant samples following the protocol by Doyle and Doyle (1987). Approximately 1 ng of DNA was used as a template for whole genome amplification via rolling circle amplification (RCA) (Inoue-Nagata et al., 2004). RCA products were digested with restriction enzymes to yield linear, genomic-length molecules (∼2,600 bp), which were cloned into the pBLUESCRIPT-KS+ plasmid vector (Agilent Technologies). Recombinant plasmids were introduced into *Escherichia coli* DH5α competent cells by electroporation (Sambrook and Russel, 2001), and the clones were fully sequenced at Macrogen, Inc. (Seoul, South Korea).

### Phylogenetic analysis

Contigs were assembled using Geneious software (Kearse et al., 2012) and oriented to start at the cleavage site of the replication origin (TAATATT//AC), following the convention established for geminiviruses (Fiallo-Olive et al., 2021). An initial BLASTn search against the GenBank database (Altschul et al., 1990) was performed to determine the viruses closest to the analyzed sequences. Taxonomic assignments were carried out by pairwise sequence comparisons using Sequence Demarcation Tool (SDT) v. 1.2, according to the guidelines of the *Geminiviridae* Study Group of the ICTV (Brown et al., 2015). Begomovirus isolates with >91% pairwise identity corresponds to the same species, and isolates from the same species that present >94% identity are considered as belonging to the same strain (Brown et al., 2015). Genetic content and genomic organization analyses were carried out using ORFfinder (Sayers et al., 2022).

Multiple sequence alignments were constructed for each of the genomic components (DNA-A and DNA-B) as well as each of the ORFs encoded by each component using the MAFFT v. 7 algorithm (Katoh and Standley, 2013), using default parameters. Maximum likelihood phylogenetic analysis was performed using RaxML-NG (Kozlov et al., 2019). The most suitable evolutionary model was determined using ModelTest-NG based on the Akaike Information Criterion (AIC) (Darriba et al., 2020). The analyses were performed using 1,000 bootstrap replicates.

### Assessment of begomovirus diversity over time

Diversity comparisons between samples collected on different years were based on the effective number of species (Hill numbers) (Hill, 1973) using the iNEXT package of R software (Hsieh et al., 2016). This metric allows the quantification of the species richness of a system, as well as their evenness. Diversity orders are estimated using the scaling parameter q (Alberdi and Gilbert, 2019). A diversity of order zero (q=0) is insensitive to the frequency of species in the container, thus favoring rare species and penalizing relative abundance. A value of q=1 weights the frequency of species and is not inflated by either rare or abundant species. A diversity of order two (q=2) is inflated by more abundant species (Alberdi and Gilbert, 2019; Jost, 2006). We defined our type as the number of DNA-A sequences of a given species per year normalized to the number of plants analyzed each year. Since the number of plants analyzed was not the same each year, we performed rarefaction and extrapolation analyses using iNEXT (Hsieh et al., 2016) to evaluate whether the sample size would have been sufficient to capture viral diversity.

In addition to the conventional approach for estimating Hill numbers, we monitored the temporal trajectory of species that make up the viral community using a recently developed approach named KHILL (Narechania et al., 2024). The basis of the tool is the same, but it differs from the previous approach in that it is not based on counting to calculate diversity. Instead, this strategy is based on the diversity of information contained in k-mer libraries obtained from genomic data. KHILL performs comparisons between unaligned substrings of length = k, thus the containers are considered as libraries of k-mers. The sets of k-mers were determined based on complete sequences, using sliding windows of k = 19 (the program’s default). betaEnt is a function of the sum of the contribution of each individual genome to total entropy, which in turn is given by the frequency at which this k-mer is represented within the sample. Total entropy is then used to determine KHILL, which is given by exp(betaEnt), providing a measure of diversity (Narechania et al., 2024). The amount of information is thus accumulated, allowing for comparisons between containers based on the KHILL curve. Very homogeneous data sets (with high information overlap) will produce a KHILL equal to 1, while complex data sets without information overlap will generate a higher KHILL number. The metric takes relative abundance into account, as populations with several species distributed in uniform proportions will produce higher KHILL values than populations where one species prevails. This method is free from sequence alignment and tree construction. As in the previously described approach, our containers were defined based on the sampling years.

### Estimates of genetic variability

We calculated the average nucleotide diversity index, π (Nei, 1987), for both genomic segments and for each gene. Estimates of π values were computed through pairwise comparisons using a Python script developed by Lima et al. (2017). We then computed a 95% confidence interval for the mean values of π through a bootstrap test with 1,000 non-parametric simulations using the boot package in R software (https://cran.r-project.org/web/packages/boot/boot.pdf). Additional variability indices were computed using DnaSP v. 6 (Rozas et al., 2017). Using phylogenetic methods, we inferred mutational biases in the complete genome datasets of each species. The number of nucleotide substitutions was estimated by mapping base changes onto the phylogenetic tree of each species using parsimony methods implemented in PAUP* (Swofford, 2003). We then compared the deviations of the observed frequencies of each substitution type with their expected values through χ² significance tests.

### Recombination analysis

To investigate the occurrence of recombination events, multiple alignments of complete DNA-A and DNA-B sequences were scanned separately using the Rdp, Geneconv, Bootscan, Maximum χ2, Chimaera, Siscan and 3Seq methods implemented in the RDP5 package (Martin et al., 2021), using default parameters. Statistical significance was inferred by P values lower than a Bonferroni-corrected α = 0.05 cutoff. As recommended by the authors, only recombination events detected by at least four methods were considered reliable.

Phylogenetic networks were built with the aim of capturing phylogenetic inconsistencies caused by hybridization. The networks were inferred using the Neighbor-Net algorithm implementing in SplitsTree v. 4.19.2 (Huson and Bryant, 2006).

To investigate recombination events on a fine scale, recombination rates were inferred throughout the genome. Watterson’s estimator of the mutation rate (θW) was calculated using the pairwise package in LDHat (McVean et al., 2002). Furthermore, we estimated the relative contribution of mutation and recombination to the diversification of the populations by calculating the population-scaled ratio between recombination and mutation rates (ρ/θW). Some adjustments needed to be done to the OxYVV data set, since the likelihood lookup table available with theta =0.001 goes up to only 120 sequences, and the OxYVV data set exceeded this value. As the computational cost to generate a likelihood lookup table is high, we adjusted the data set using CD-Hit (Fu et al., 2012), eliminating sequences with >99.96% identity and thus reducing the data set to 99 sequences.

### Temporal signal evaluation and demographic analysis

The temporal signal strength was evaluated using Bayesian Evaluation of Temporal Signal (BETS) (Duchene et al., 2020) implemented in BEAST v1.10.4. Two scenarios were considered. In the first one the data was treated as being isochronous. In the second one we treated the data as heterochronous (Suchard et al., 2018), estimated the marginal likelihood using generalized stepping-stone sampling (GSS) (Baele et al., 2015), and then computed the Bayes factor (Kass and Raftery, 1995). The molecular clock that best suited the data was also determined based on the Bayes factor (Kass and Raftery, 1995), with an MCMC chain length of 2 x 10^8^ with 10% burn in, and a number of stepping stones of 100, with an MCMC length of 2 × 10^6^. The convergence of the analysis was assessed using Tracer v. 1.7, and was determined based on the effective sample size, which should assume values of at least 200, and by the degree of interdependence of the samples assessed based on the degree of mixing of the parameters (Rambaut et al., 2018). The time scale trees were summarized using the Maximum Clade Credibility (MCC) method in TreeAnnotaor v2.6.4 (Drummond and Rambaut, 2007). The substitution/site/year rates and Time to the Most Recent Common Ancestor (TMRCA) were estimated with a 95% HPD confidence interval. The trajectory of populations over time was monitored through the construction of Bayesian Skygrid coalescent models (Hill and Baele, 2019).

### Selection analysis

Selection analysis was performed for each of the coding regions of the genome of each species to identify potential sites that could be experiencing positive or negative selection, using three methods: Single Likelihood Ancestor Counting (SLAC) (Kosakovsky-Pond and Frost, 2005), Mixed Effects Model of Evolution (MEME) (Murrell et al., 2012) and Fast Unconstrained Bayesian AppRoximation (FUBAR) (Murrell et al., 2013), all implemented in the DataMonkey webserver (Weaver et al., 2018). Relationships between synonymous and non-synonymous substitution rates (ω) were calculated using SLAC. To avoid any artifacts in our analyses, a search for recombination breakpoints was carried out for each data set using the Genetic Algorithm Recombination Detection (GARD) (Pond et al., 2006). We also performed Tajima’s D neutrality tests and calculated Fu and Li’s D* and F* statistics in DnaSP v. 6 (Rozas et al., 2017).

## Results

### *Sida acuta* is a “mixing vessel” host for begomoviruses

Over the course of 12 years, we collected 360 *Sida acuta* plants with symptoms of begomovirus infection (Suppl. Figure S1) in the same small area (300 m^2^) near Viçosa, MG (Suppl. Figure S2), On average, 35 plants were collected per year. A total of 371 clones corresponding to full-length begomovirus components were obtained from 242 individual plant samples, 262 corresponding to a DNA-A and 109 corresponding to a DNA-B (Suppl. Table S1).

Based on complete DNA-A and DNA-B sequences, begomoviruses belonging to five different species were detected in the area (Suppl. Table S1; Suppl. Table S2). The most abundant was Oxalis yellow vein virus (OxYVV; *Begomovirus oxalisflavi*), from which we obtained a total of 162 DNA-A (127 haplotypes) and 52 DNA-B clones (40 haplotypes), followed by Sida yellow leaf curl virus (SiYLCV; *Begomovirus sidaflavacontorsionis*), which accounted for 77 DNA-A (59 haplotypes) and 44 DNA-B clones (35 haplotypes), Sida micrantha mosaic virus (SimMV; *Begomovirus sidamicranthae*) with 17 DNA-A (10 haplotypes) and 9 DNA-B (6 haplotypes) clones, Sida mottle virus (SiMV; *Begomovirus sidavariati*) with 5 DNA-A and 4 DNA-B clones, and a single DNA-A clone of Macroptilium yellow vein virus (MaYVV; *Begomovirus macroptilivenae*).

Comparison of the DNA-A sequences indicated that the OxYVV population comprises two distinct strains, S1 and S2, with isolates from each strain sharing a maximum DNA-A identity of 93.8% with those from the other strain (Suppl. Figure S3). The S1 strain is much more prevalent, totaling 155 clones, and can be subdivided into five distinct groups (S1a-e), which we named variants since they present pairwise identity percentages above the strain threshold (95-97% among variants; Suppl. Figure S3). The remaining seven OxYVV clones correspond to the S2 strain. For SiYLCV, the population is comprised of a single variant, with 97.5-98% identities among all clones (Suppl. Figure S4). The SimMV population harbors two highly divergent variants which are very close to the strain demarcation threshold (93.8-94.5% identity between clones of the two variants; Suppl. Figure S5).

The phylogenetic tree inferred from the complete DNA-A sequences showed four well supported clades and one long branch, each corresponding to one of the species mentioned above (Figure 1A). The clade corresponding to OxYVV branches into six subclades corresponding to the five variants of strain S1 and the one group of strain S2 sequences. The bootstrap values of some of the branches that separate the S1 variants are low, possibly due to the high identity values between individual sequences. The clade that harbors SiYLCV also presents several subclades but with short genetic distances, consistent with the classification of all the isolates as a single variant (Figure 1A).

**Figure 1.**
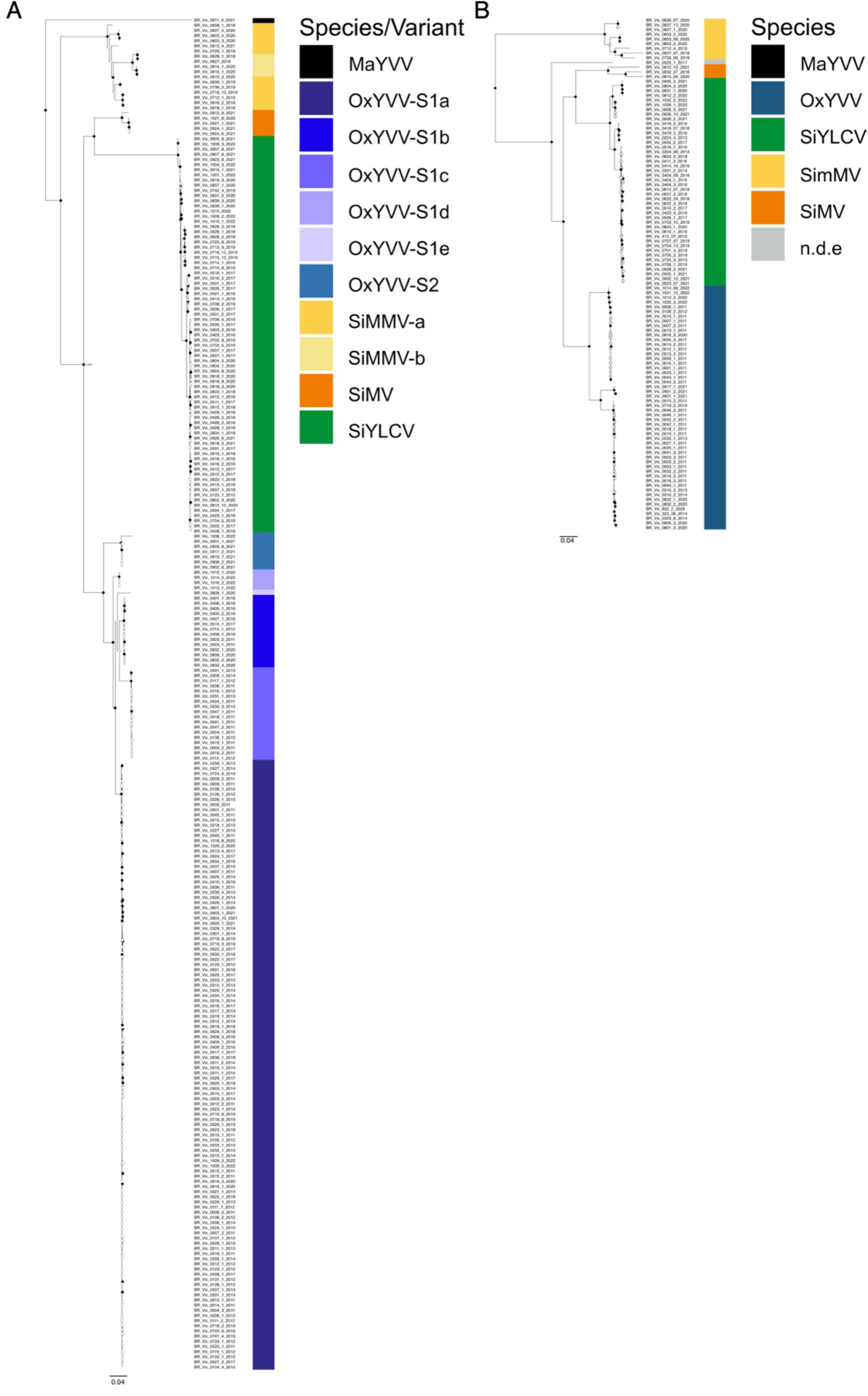
Phylogenetic trees based on the complete DNA-A **(A)** and DNA-B **(B)** nucleotide sequences of the begomovirus isolates obtained in this work. Nodes to the right of branches with bootstrap support (1000 replications) equal to or higher than 75% are indicated by black circles, and those with values lower than 75% by white circles. Isolates classified as members of the species Oxalis yellow vein virus (OxYVV) are represented in blue (or different shades of blue for the DNA-A tree), Sida yellow leaf curl virus (SiYLCV) in green, Sida micrantha mosaic virus (SimMV) in yellow, Sida motlle virus (SiMV) in orange, and Macroptilium yellow vein virus (MaYVV) in black. An isolate for which species assignment was not possible is represented in grey in the DNA-B tree. Species, strains and variants were identified based on the demarcation criteria established by the *Geminiviridae* Study Group of the ICTV (Brown et al., 2015). The scale bar represents nucleotide substitutions per site.

The phylogenetic tree based on the complete DNA-B sequences also formed four large clades corresponding to the four most prevalent species (Figure 1B). The clade corresponding to OxYVV is subdivided into three well-supported subclades. A second clade corresponding to SiYLCV is subdivided into four subclades, all well supported, but with shorter genetic distances when compared to the subclades formed by OxYVV isolates (Figure 1B). The tree also has a long, distinctive branch corresponding to a DNA-B sequence that probably represents a new species. However, since a cognate DNA-A (on which taxonomy is based) (Brown et al., 2015) was not cloned, it is not possible to confirm the identity of this isolate.

Together, these results indicate that the *Sida acuta* plants harbor a complex begomovirus community composed of at least five viruses, some of which can be divided into strains and variants. Taking into consideration the small sampling area, these results demonstrate how non-cultivated hosts may constitute sources of viral diversity.

### Temporal dynamics reveal fluctuations in the frequencies of viruses and their variants in the landscape

The temporal dynamics of begomoviruses in the community were characterized based on the distribution of the relative frequencies of DNA-A for each species, strain, and variant, given that this genomic component has taxonomic value and underpins the definition of these groups. Changes were observed in the composition of species and variants over the years (Figure 2; Suppl. Table S2). OxYVV was the prevalent virus from 2011 to 2014, with three variants present (OxYVV-S1a, -S1b, -S1c). OxYVV-S1a was at a higher frequency since the first year and its frequency increased until 2014. The OxYVV-S1b variant had a lower frequency during this period, not being detected in 2013 or 2014. The OxYVV-S1c variant appeared in second place in terms of frequency, but its incidence was reduced in subsequent years. In 2013, a single plant infected with SiYLCV was detected for the first time. The year 2015 was a gap in our sampling. In 2016 there was a major change in the proportion of species and variants present at the site, with SiYLCV outnumbering OxYVV as the prevalent virus. Another change was that the OxYVV-S1b variant was detected again and this time at a higher frequency than OxYVV-S1a. In 2017 the frequency of variant OxYVV-S1a increased again, while OxYVV-S1b and SiLCV had a slight reduction in their frequencies. In 2018 we observed the local emergence of SimMV, with OxYVV-S1 and SiYLCV exhibiting balanced frequencies between themselves. In the three subsequent years there were fluctuations in the frequencies of these three viruses, but always with an alternation between OxYVV and SiYLCV in terms of prevalence. The variant OxYVV-S1e emerged in 2020. The year 2021 marked the first detection of MaYVV and SiMV, in addition to the OxYVV-S2 strain. In 2022 we detected the two strains of OxYVV and the emergence of OxYVV-S1d (Figure 2B).

**Figure 2.**
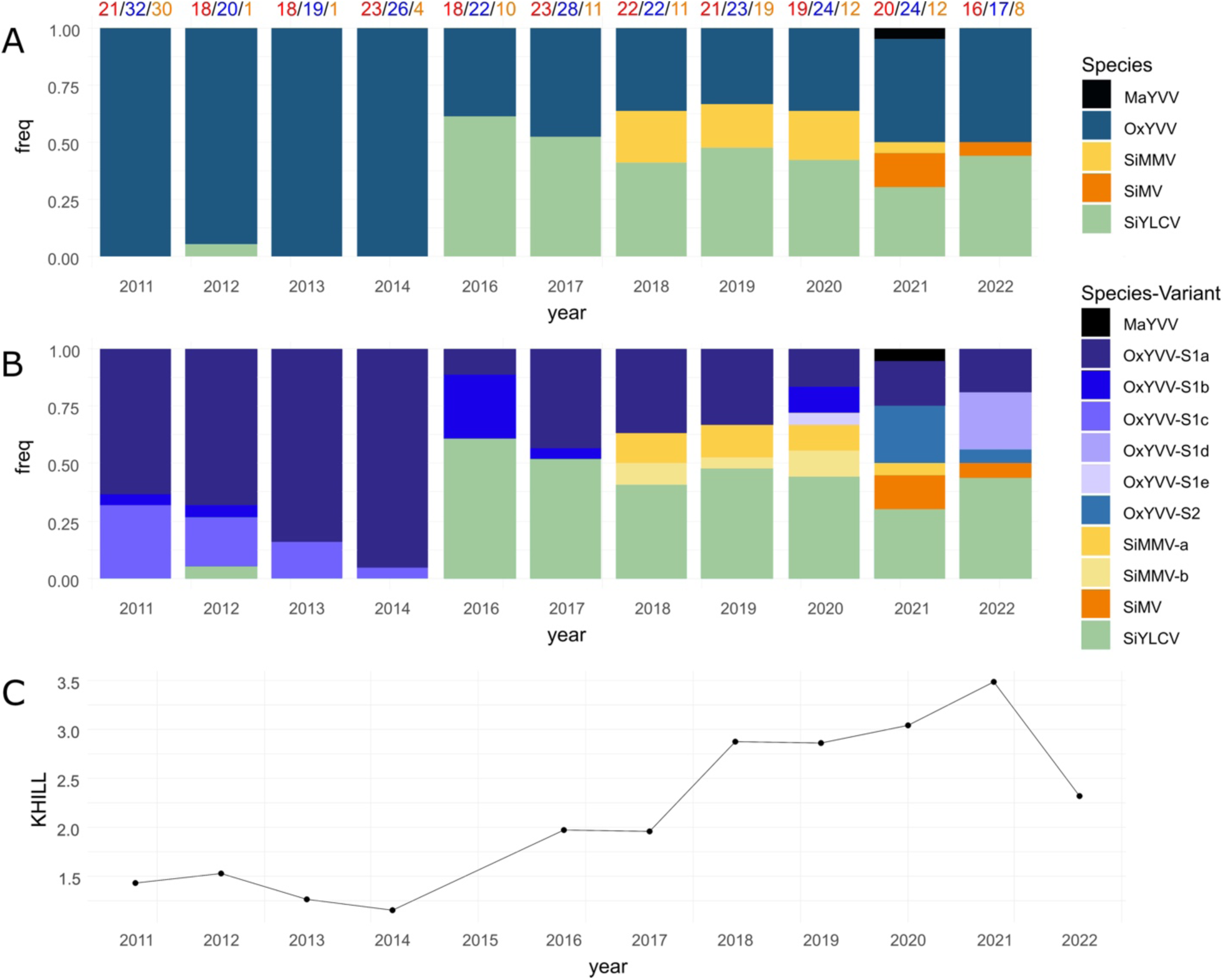
**A.** Proportion of individuals of each begomovirus sampled in each year. The data is based on the sum of DNA-A and DNA-B sequences obtained each year. The color scale on the right indicates species, strain and variant assignments. The numbers at the top of the bars indicate the number of plants analyzed (red), the number of DNA-A sequences recovered (blue), the number of DNA-B sequenced (orange). MaYVV, Macroptilium yellow vein virus; OxYVV, Oxalis yellow vein virus; SiYLCV, Sida yellow leaf curl virus; SimMV, Sida micrantha mosaic virus; SiMV, Sida motlle virus. **B.** Relative proportion of each species and their respective variants isolated in each year. The data is based on DNA-A sequences only, since there are no criteria for strain and variant assignment based on the DNA-B. **C.** KHILL values, which are estimated based on the entropy and represents the dynamics of the landscape over the years.

Hill numbers were calculated to assess diversity over time. We adopted this strategy because it translates into a simple and intuitive measure that allows us to compare the diversity of species that make up a community. We opted to use species counts to assess diversity; another option would be to use variant diversity, but this could make our interpretation confusing (see below). The years 2011, 2013 and 2014 had only one species detected (Figure 2A), in which case the value converged to unity (Suppl. Figure S6). The years 2012, 2016 and 2017 also had a very low diversity, and the overlap of the three orders of diversity in 2016 and 2017 indicates that two species were represented in similar proportions in those years (Suppl. Figure S6). In the years 2018, 2019, 2020 and 2022 the richness values were equal to 3, and in 2021 it was equal to 5, the highest value observed in our time series.

There was no significant difference between diversity orders 1 and 2 from 2018 to 2022 (Suppl. Figure S6; Suppl. Table S1). The effective number of species approached richness in the years 2016, 2017, 2018 e 2020, indicating that in these years the community was dominated by equally abundant species.

To capture variations in diversity on a finer scale, we used KHILL, an approach which is also based on Hill numbers, but which uses DNA sequences and calculates the degree of entropy. We noticed changes in the behavior of the KHILL curve that follow the dynamics of diversity changes over time (Figure 2C). The KHILL curve was able to effectively capture changes in the diversity of species present at the site and beyond, detecting nuances in the dynamics of variants within the community. This can be observed in the first four years, as the KHILL values decrease together with the reduction in incidence of minor variants of OxYVV (Figure 2C). This was followed by a sudden increase starting in 2016 until it peaked in 2021, when five species were detected.

### Measuring the genetic variability of begomoviruses in *Sida acuta*

The genetic variability of each virus was inferred separately. Due to the size of the data sets, we did these estimates only for three of the five viruses detected in the area (OxYVV, SiYLCV SimMV). The other two viruses (SiMV and MaYVV) were represented by a very small number of sequences (5 and 1, respectively). We estimated the variability for each genomic component and also for their genes by calculating the nucleotide diversity index, π (Table 1). In general, the DNA-B showed greater genetic variability compared to the DNA-A, as has already been reported for most bipartite begomoviruses (Briddon et al., 2010; Xavier et al., 2021). For both components, SimMV was the virus that showed the greatest variability, followed by OxYVV and then SiYLCV (Table 1). When we looked at each gene separately, *Rep* and *Ren* were the ones that showed the greatest variability. For the DNA-B, the *MP* gene of SimMV and OxYVV showed greater variability, while for SiYLCV the *NSP* gene had higher variability (Table 1).

**Table 1.**
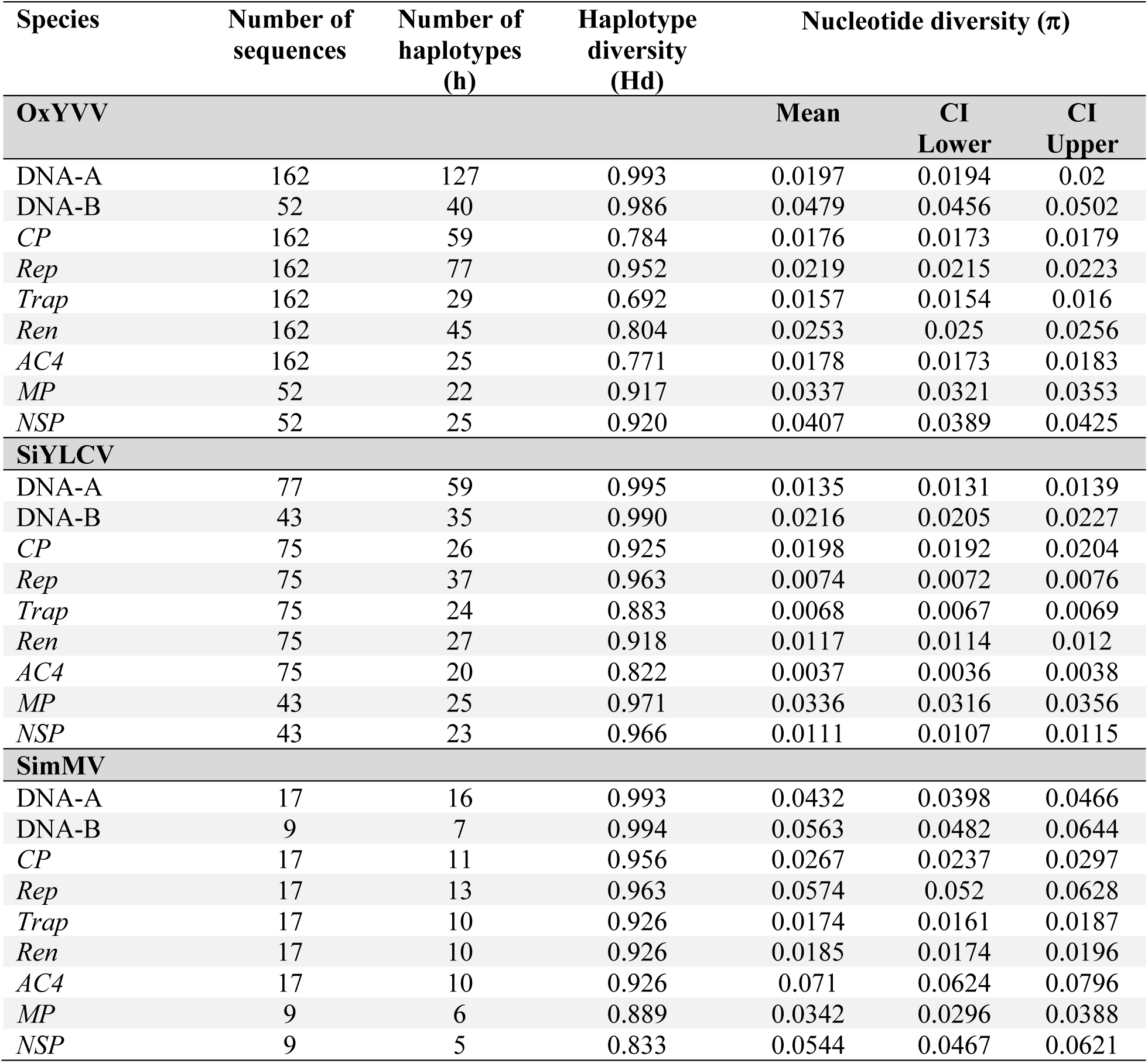
Genetic variability indices of the begomoviruses infecting *Sida acuta* in Viçosa, MG.

Although all viruses were sampled from the same host species, large differences in nucleotide diversity were observed among them. To assess whether specific mechanisms were favoring certain types of mutations in the viral genomes, we estimated mutational bias based on base frequencies. We detected multiple significant biases, with a consistent trend across the three species and both genomic components. Among these, C→T and G→A substitutions showed the strongest deviations according to the χ² test (Table 2). These patterns are widely reported in begomoviruses (Duffy and Holmes, 2008; Duffy and Holmes, 2009; van der Walt et al., 2008) and can be largely explained by cytosine deamination to uracil (read as thymine during replication), adenine deamination to hypoxanthine (which pairs as guanine), and guanine deamination to xanthine (which pairs as thymine, ultimately leading to G→A transitions), respectively (Caulfield et al., 1998). Interestingly, the DNA-B of OxYVV exhibited a significant excess of G→T substitutions, a mutation type commonly linked to oxidative guanine damage. This bias is consistent with the nature of single-stranded DNA genomes, in which exposed bases are more prone to guanine oxidation, generating 8-oxoG that can mispair with adenine during replication (Hahm et al., 2022; van der Walt et al., 2008; Wood et al., 1992).

**Table 2.** Mutational biases inferred from the maximum likelihood phylogeny of Oxalys yellow vein virus (OxYVV), Sida yellow leaf curl virus (SiYLCV), and Sida micrantha mosaic virus (SimMV) for both DNA-A and DNA-B components, evaluated using χ² tests of significance. Each cell shows, in order, the number of observed substitutions, the expected number of substitutions (in parentheses), the χ² statistic, and the corresponding *P*-value.

| OxYVV- DNA-A |  |  |  |  |
| --- | --- | --- | --- | --- |
| Resulting base |  |  |  |  |
| Original base | A | T | C | G |
| A | - | 35 (18.5), 14.716, 0.00012 | 21 (13.2), 4.533, 0.033 | 71.5 (38.0), 29.533, 5.5e<sup>-08</sup> |
| T | 39 (18.5), 22.716, 1.9e<sup>-06</sup> | - | 121.5 (71.5), 34.965, 3.4e<sup>-09</sup> | 36 (32.0), 0.500, 0.48 |
| C | 32 (13.2), 26.533, 2.6e<sup>-07</sup> | 164.5 (71.5), 120.965, 3.9e<sup>-28</sup> | - | 40 (19.5), 21.551, 3.4e<sup>-06</sup> |
| G | 80.5 (38.0), 47.533, 5.4e<sup>-12</sup> | 92 (32.0), 112.500, 2.8e<sup>-26</sup> | 38 (19.5), 17.551, 2.8e<sup>-05</sup> | - |
| OxYVV- DNA-B |  |  |  |  |
| Resulting base |  |  |  |  |
| Original base | A | T | C | G |
| A | - | 31 (15.5), 15.500, 8.3e<sup>-05</sup> | 16 (7.8), 8.782, 0.003 | 54 (27.2), 26.259, 3e<sup>-07</sup> |
| T | 31 (15.5), 15.500, 8.3e<sup>-05</sup> | - | 65 (37.0), 21.189, 4.2e<sup>-06</sup> | 43 (22.8), 18.025, 2.2e<sup>-05</sup> |
| C | 15 (7.8), 6.782, 0.0092 | 83 (37.0), 57.189, 4e<sup>-14</sup> | - | 18 (8.8), 9.779, 0.0018 |
| G | 55 (27.2), 28.259, 1.1e<sup>-07</sup> | 48 (22.8), 28.025, 1.2e<sup>-07</sup> | 17 (8.8), 7.779, 0.0053 | - |
| SiYLCV-DNA-A |  |  |  |  |
| Resulting base |  |  |  |  |
| Original base | A | T | C | G |
| A | - | 7 (5.0), 0.800, 0.37 | 6 (3.5), 1.786, 0.18 | 25 (14.2), 8.110, 0.0044 |
| T | 13 (5.0), 12.800, 0.00035 | - | 37.5 (27.8), 3.426, 0.064 | 23 (12.2), 9.434, 0.0021 |
| C | 8 (3.5), 5.786, 0.016 | 73.5 (27.8), 75.426, 3.8e<sup>-18</sup> | - | 8 (4.0), 4.000, 0.046 |
| G | 32 (14.2), 22.110, 2.6e<sup>-06</sup> | 26 (12.2), 15.434, 8.5e<sup>-05</sup> | 8 (4.0), 4.000, 0.046 | - |
| SiYLCV-DNA-B |  |  |  |  |
| Resulting base |  |  |  |  |
| Original base | A | T | C | G |
| A | - | 20 (10.8), 7.959, 0.0048 | 12 (6.8), 4.083, 0.043 | 30 (16.0), 12.250, 0.00047 |
| T | 23 (10.8), 13.959, 0.00019 | - | 50.5 (29.0), 15.940, 6.5e<sup>-05</sup> | 20 (12.2), 4.903, 0.027 |
| C | 15 (6.8), 10.083, 0.0015 | 65.5 (29.0), 45.940, 1.2e<sup>-11</sup> | - | 24 (9.5), 22.132, 2.5e<sup>-06</sup> |
| G | 34 (16.0), 20.250, 6.8e<sup>-06</sup> | 29 (12.2), 22.903, 1.7e<sup>-06</sup> | 14 (9.5), 2.132, 0.14 | - |

**Table 2** (cont.)
| SimMV- DNA-A |  |  |  |  |
| --- | --- | --- | --- | --- |
| Resulting base |  |  |  |  |
| Original base | A | T | C | G |
| A | - | 25.5 (11.2), 18.050 2.2e<sup>-05</sup> | 18 (7.8), 13.556, 0.00023 | 36.5 (22.8), 8.310, 0.039 |
| T | 19.5 (11.2), 6.050, 0.014 | - | 82 (42.2), 37.398, 9.6e<sup>-10</sup> | 37.5 (19.2), 17.302, 3.2e<sup>-05</sup> |
| C | 13 (7.8), 3.556, 0.059 | 87 (42.2), 47.398, 5.8e<sup>-12</sup> | - | 17 (8.0), 10.125, 0.0015 |
| G | 54.5 (22.8), 44.310, 2.8e<sup>-11</sup> | 39.5 (19.2), 21.302, 3.9e<sup>-06</sup> | 15 (8.0), 6.125, 0.013 | - |
| SimMV- DNA-B |  |  |  |  |
| Resulting base |  |  |  |  |
| Original base | A | T | C | G |
| A | - | 41 (18.8), 26.403, 2.8e<sup>-07</sup> | 20 (10.0), 10.000, 0.0016 | 48 (24.2), 23.260, 1.4e<sup>-06</sup> |
| T | 34 (18.8), 12.403, 0.00043 | - | 70 (36.0), 32.111, 1.5e<sup>-08</sup> | 39 (21.0), 15.429, 8.6e<sup>-05</sup> |
| C | 20 (10.0), 10.000, 0.0016 | 74 (36.0), 40.111, 2.4e<sup>-10</sup> | - | 30 (12.2), 25.719, 3.9e<sup>-07</sup> |
| G | 49 (24.2), 25.260, 5e<sup>-07</sup> | 45 (21.0), 27.429, 1.6e<sup>-07</sup> | 19 (12.2), 3.719, 0.054 | - |

### Recombination analysis

Since the begomoviruses sampled in this study are part of the same community (with several cases of mixed infections; Suppl. Table S1), recombination was analyzed in an inter-specific data set including sequences from the five viruses. Recombination events involving the DNA-A of at least one individual from each sampled species were identified (Table 3). Only one recombination event was detected for OxYVV, in an isolate (BR_VIC1006_1) of the S2 strain. The event involves a major parent of the same strain and an unknown minor parent. Interestingly, all 77 SiYLCV sequences were identified as recombinants, in an event involving OxYVV as a minor parent and an unknown major parent. Three recombination events were detected for SimMV, one of which was strictly intraspecific and the other two involved minor parents of the same species and unknown major parents. Two events were detected for SiMV, the first involving OxYVV and SiYLCV and the second involving SiYLCV and SimMV. We also detected evidence of recombination in MaYVV involving OxYVV as the main parent. Most of the recombination breakpoints were located between the intergenic region and the *Rep* gene, the only exception being event 8 where the breakpoints were located in the *CP* gene (Table 3).

**Table 3.**
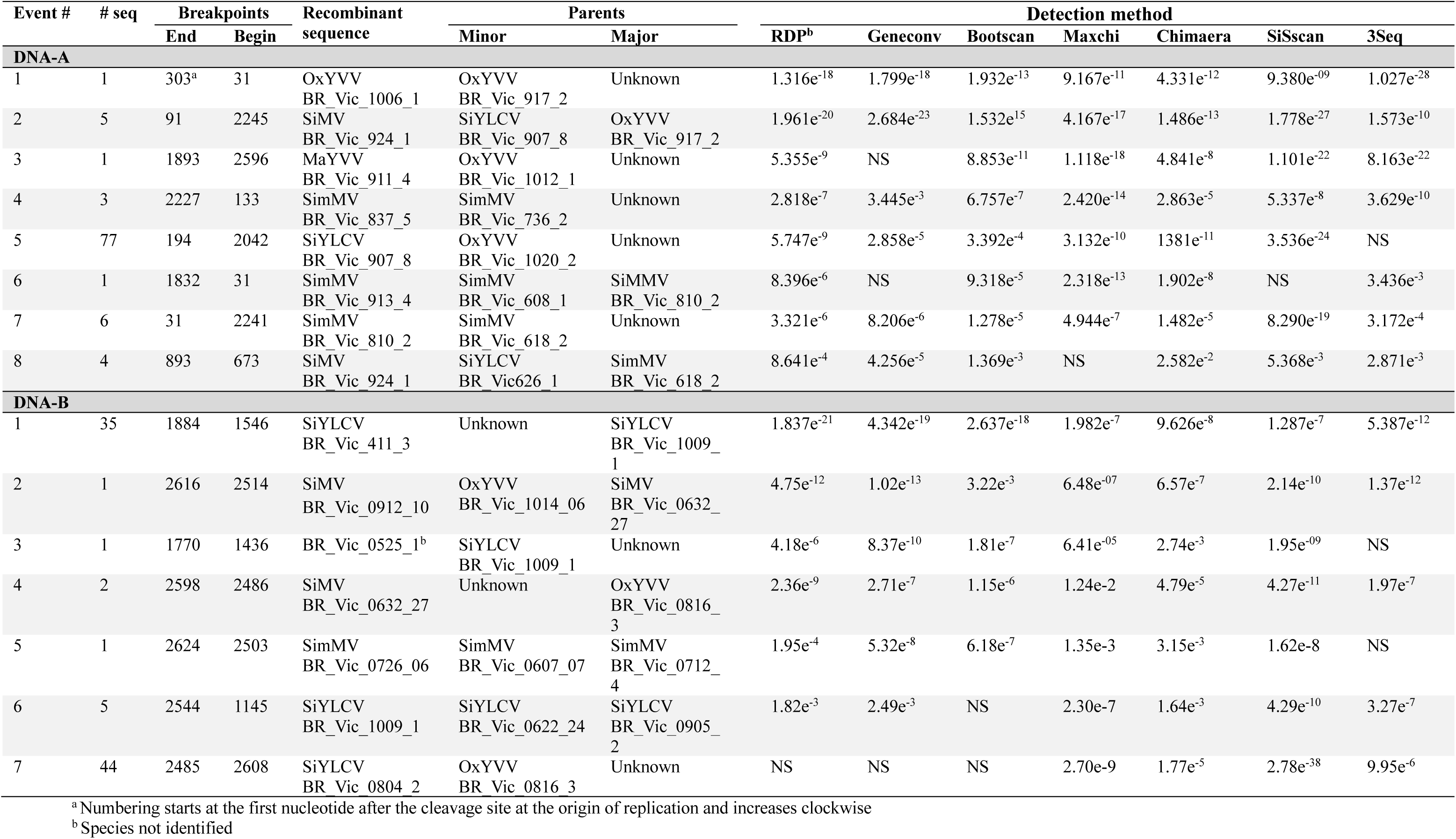
Summary of recombination events detected in the begomovirus community infecting *Sida acuta* in Viçosa, MG. The analysis was performed using complete DNA-A and DNA-B sequences.

| Event # | # seq | Breakpoints |  | Recombinant sequence | Parents |  | Detection method |  |  |  |  |  |  |
| --- | --- | --- | --- | --- | --- | --- | --- | --- | --- | --- | --- | --- | --- |
|  |  | End | Begin |  | Minor | Major | RDP<sup>b</sup> | Geneconv | Bootscan | Maxchi | Chimaera | SiSscan | 3Seq |
| DNA-A |  |  |  |  |  |  |  |  |  |  |  |  |  |
| 1 | 1 | 303<sup>a</sup> | 31 | OxYVV<br>BR_Vic_1006_1 | OxYVV<br>BR_Vic_917_2 | Unknown | 1.316e<sup>-18</sup> | 1.799e<sup>-18</sup> | 1.932e<sup>-13</sup> | 9.167e<sup>-11</sup> | 4.331e<sup>-12</sup> | 9.380e<sup>-09</sup> | 1.027e<sup>-28</sup> |
| 2 | 5 | 91 | 2245 | SiMV<br>BR_Vic_924_1 | SiYLCV<br>BR_Vic_907_8 | OxYVV<br>BR_Vic_917_2 | 1.961e<sup>-20</sup> | 2.684e<sup>-23</sup> | 1.532e<sup>15</sup> | 4.167e<sup>-17</sup> | 1.486e<sup>-13</sup> | 1.778e<sup>-27</sup> | 1.573e<sup>-10</sup> |
| 3 | 1 | 1893 | 2596 | MaYVV<br>BR_Vic_911_4 | OxYVV<br>BR_Vic_1012_1 | Unknown | 5.355e<sup>-9</sup> | NS | 8.853e<sup>-11</sup> | 1.118e<sup>-18</sup> | 4.841e<sup>-8</sup> | 1.101e<sup>-22</sup> | 8.163e<sup>-22</sup> |
| 4 | 3 | 2227 | 133 | SimMV<br>BR_Vic_837_5 | SimMV<br>BR_Vic_736_2 | Unknown | 2.818e<sup>-7</sup> | 3.445e<sup>-3</sup> | 6.757e<sup>-7</sup> | 2.420e<sup>-14</sup> | 2.863e<sup>-5</sup> | 5.337e<sup>-8</sup> | 3.629e<sup>-10</sup> |
| 5 | 77 | 194 | 2042 | SiYLCV<br>BR_Vic_907_8 | OxYVV<br>BR_Vic_1020_2 | Unknown | 5.747e<sup>-9</sup> | 2.858e<sup>-5</sup> | 3.392e<sup>-4</sup> | 3.132e<sup>-10</sup> | 1381e<sup>-11</sup> | 3.536e<sup>-24</sup> | NS |
| 6 | 1 | 1832 | 31 | SimMV<br>BR_Vic_913_4 | SimMV<br>BR_Vic_608_1 | SiMMV<br>BR_Vic_810_2 | 8.396e<sup>-6</sup> | NS | 9.318e<sup>-5</sup> | 2.318e<sup>-13</sup> | 1.902e<sup>-8</sup> | NS | 3.436e<sup>-3</sup> |
| 7 | 6 | 31 | 2241 | SimMV<br>BR_Vic_810_2 | SimMV<br>BR_Vic_618_2 | Unknown | 3.321e<sup>-6</sup> | 8.206e<sup>-6</sup> | 1.278e<sup>-5</sup> | 4.944e<sup>-7</sup> | 1.482e<sup>-5</sup> | 8.290e<sup>-19</sup> | 3.172e<sup>-4</sup> |
| 8 | 4 | 893 | 673 | SiMV<br>BR_Vic_924_1 | SiYLCV<br>BR_Vic626_1 | SimMV<br>BR_Vic_618_2 | 8.641e<sup>-4</sup> | 4.256e<sup>-5</sup> | 1.369e<sup>-3</sup> | NS | 2.582e<sup>-2</sup> | 5.368e<sup>-3</sup> | 2.871e<sup>-3</sup> |
| DNA-B |  |  |  |  |  |  |  |  |  |  |  |  |  |
| 1 | 35 | 1884 | 1546 | SiYLCV<br>BR_Vic_411_3 | Unknown | SiYLCV<br>BR_Vic_1009_1 | 1.837e<sup>-21</sup> | 4.342e<sup>-19</sup> | 2.637e<sup>-18</sup> | 1.982e<sup>-7</sup> | 9.626e<sup>-8</sup> | 1.287e<sup>-7</sup> | 5.387e<sup>-12</sup> |
| 2 | 1 | 2616 | 2514 | SiMV<br>BR_Vic_0912_10 | OxYVV<br>BR_Vic_1014_06 | SiMV<br>BR_Vic_0632_27 | 4.75e<sup>-12</sup> | 1.02e<sup>-13</sup> | 3.22e<sup>-3</sup> | 6.48e<sup>-07</sup> | 6.57e<sup>-7</sup> | 2.14e<sup>-10</sup> | 1.37e<sup>-12</sup> |
| 3 | 1 | 1770 | 1436 | BR_Vic_0525_1<sup>b</sup> | SiYLCV<br>BR_Vic_1009_1 | Unknown | 4.18e<sup>-6</sup> | 8.37e<sup>-10</sup> | 1.81e<sup>-7</sup> | 6.41e<sup>-05</sup> | 2.74e<sup>-3</sup> | 1.95e<sup>-09</sup> | NS |
| 4 | 2 | 2598 | 2486 | SiMV<br>BR_Vic_0632_27 | Unknown | OxYVV<br>BR_Vic_0816_3 | 2.36e<sup>-9</sup> | 2.71e<sup>-7</sup> | 1.15e<sup>-6</sup> | 1.24e-2 | 4.79e<sup>-5</sup> | 4.27e<sup>-11</sup> | 1.97e<sup>-7</sup> |
| 5 | 1 | 2624 | 2503 | SimMV<br>BR_Vic_0726_06 | SimMV<br>BR_Vic_0607_07 | SimMV<br>BR_Vic_0712_4 | 1.95e<sup>-4</sup> | 5.32e<sup>-8</sup> | 6.18e<sup>-7</sup> | 1.35e-3 | 3.15e<sup>-3</sup> | 1.62e-8 | NS |
| 6 | 5 | 2544 | 1145 | SiYLCV<br>BR_Vic_1009_1 | SiYLCV<br>BR_Vic_0622_24 | SiYLCV<br>BR_Vic_0905_2 | 1.82e<sup>-3</sup> | 2.49e<sup>-3</sup> | NS | 2.30e-7 | 1.64e<sup>-3</sup> | 4.29e<sup>-10</sup> | 3.27e<sup>-7</sup> |
| 7 | 44 | 2485 | 2608 | SiYLCV<br>BR_Vic_0804_2 | OxYVV<br>BR_Vic_0816_3 | Unknown | NS | NS | NS | 2.70e-9 | 1.77e<sup>-5</sup> | 2.78e<sup>-38</sup> | 9.95e<sup>-6</sup> |

<sup>a</sup>Numbering starts at the first nucleotide after the cleavage site at the origin of replication and increases clockwise
<sup>b</sup>Species not identified

For the DNA-B component, no significant recombination events were identified in the OxYVV population (Table 3). Conversely, three recombination events were detected for SiYLCV. Event 7 was detected in all SiYLCV sequences analyzed and involved OxYVV and an unknown major parent. The second event was observed in 35 SiYLCV sequences, with the major parent unknown and the minor parent being SiYLCV. The third event was an intraspecific recombination event detected in five SiYLCV sequences. A single SimMV DNA-B sequence showed evidence of intraspecific recombination. The breakpoints detected in the DNA-B varied in their location (Table 3). For four of the seven events (events 2, 4, 5 and 7) the breakpoints were found in the larger intergenic region (LIR). The breakpoints of events 1 and 3 are located within the *MP* gene, while those of event 5 spanned part of the *NSP* gene up to the LIR.

The evolutionary history and the impact of recombination were captured by reconstructing phylogenetic networks. These networks, unlike phylogenetic trees, allow us to visualize the occurrence of lateral transfers of genomic regions between individuals. The DNA-A network formed five large clusters corresponding to each of the five viruses (Figure 3A). The OxYVV cluster showed a clear division into six dense clusters with little cross-linking between them. One of these cross-links connects the BR_Vic_1006_1 isolate (OxYVV-S2 strain) to the other individuals affiliated with this strain, indicating a pattern of genetic exchange. SiYLCV formed a group separated by a long branch with rare reticulations. Some reticulations connected the SiMV and OxYVV-S2 groups, indicating patterns of recombination between these viruses. Some edges are shared within the SimMV cluster. Also, exchange patterns were found between SimMV and MaYVV.

**Figure 3.**
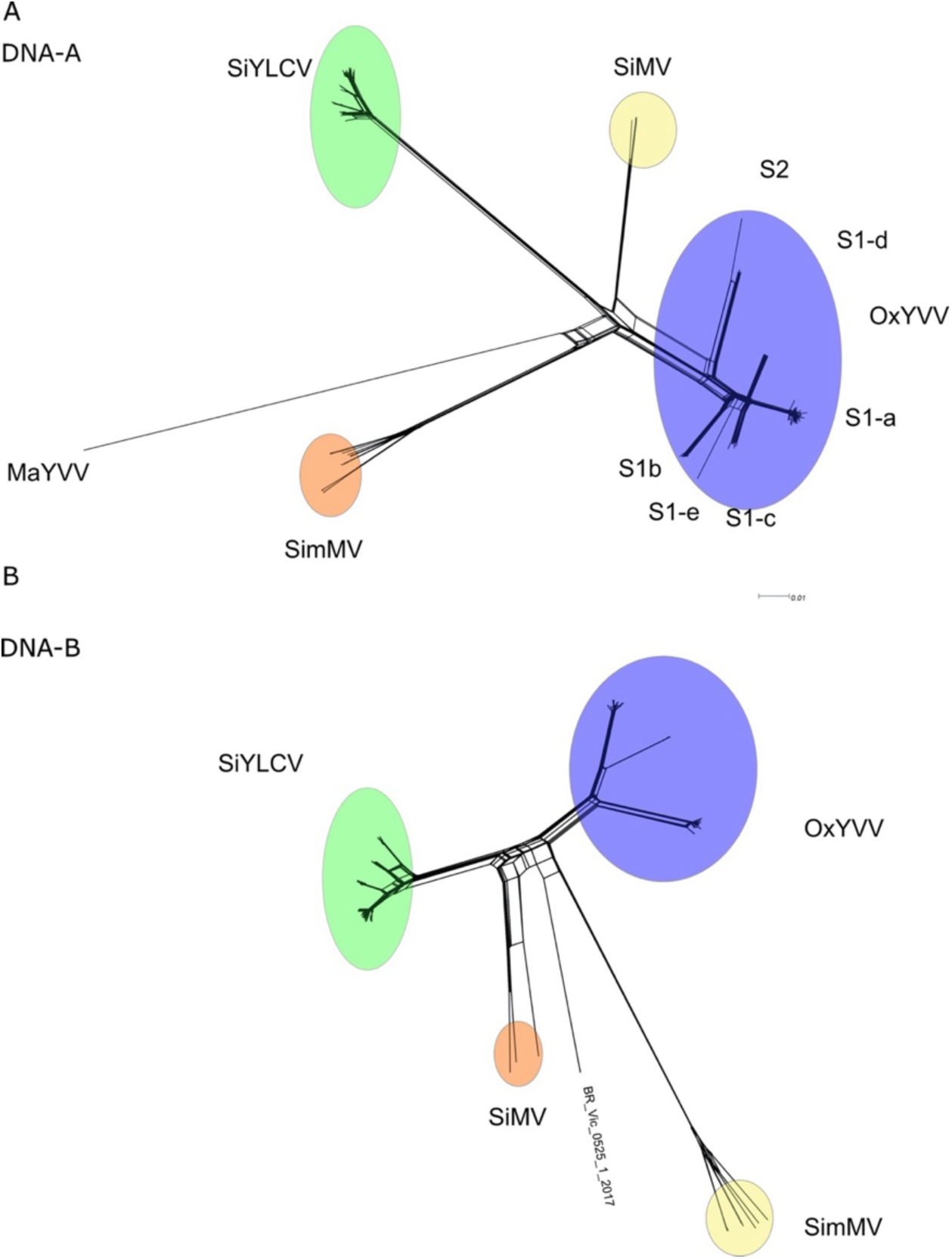
Phylogenetic evidence of recombination in the DNA-A **(A)** and DNA-B **(B)** sequences of the begomovirus community infecting *Sida acuta* in Viçosa, MG. Neighbor-Net network analysis was performed using SplitsTree4. The formation of a reticular network rather than a single bifurcated tree suggests recombination. MaYVV, Macroptilium yellow vein virus; OxYVV, Oxalis yellow vein virus; SiYLCV, Sida yellow leaf curl virus; SimMV, Sida micrantha mosaic virus; SiMV, Sida motlle virus.

The DNA-B network also presented cross-linking patterns. Five clusters were formed corresponding to each of the viruses (including the one for which the species was not identified). Edges connecting SiMV individuals were observed. Some side branches also connected SimMV and the unidentified virus, indicating possible recombination events (Figure 3B).

We inferred the relative contribution of recombination and mutation to the diversification of the *S. acuta* begomovirus community by calculating the ratio between these two rates (ρ/θ). For both components and most data sets, the ρ/θ ratio was <1 (Table 4), indicating a greater probability of polymorphisms being the result of mutation rather than recombination. The only case where there was a greater relative contribution of recombination was for SiYLCV (Table 4). Even though most of the ratios were <1, they can be considered high, revealing the important role of recombination as a driver of diversification.

**Table 4.**
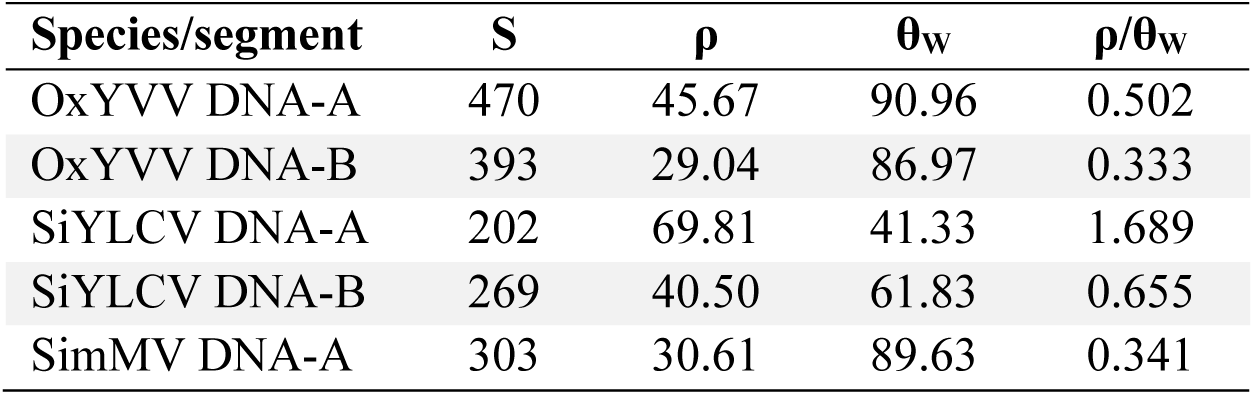
Population-scaled rates of recombination and mutation of the begomoviruses infecting *Sida acuta* in Viçosa, MG. S, number of segregating sites; ρ, population-scaled recombination rate; θw, Watterson’s estimator of population-scaled mutation rate (θ); ρ/θ, recombination/mutation ratio.

| Species/segment | S | $\rho$ | $\theta_w$ | $\rho/\theta_w$ |
| --- | --- | --- | --- | --- |
| OxYVV DNA-A | 470 | 45.67 | 90.96 | 0.502 |
| OxYVV DNA-B | 393 | 29.04 | 86.97 | 0.333 |
| SiYLCV DNA-A | 202 | 69.81 | 41.33 | 1.689 |
| SiYLCV DNA-B | 269 | 40.50 | 61.83 | 0.655 |
| SimMV DNA-A | 303 | 30.61 | 89.63 | 0.341 |

### Evidence of temporal signal in the OxYVV and SiYCV populations

We employed Bayesian Evaluation of Temporal Signal (BETS) to infer evidence of a temporal signal in the data sets. For both OxYVV and SiYLCV, the higher marginal likelihood values were observed under the heterochronous model. A Bayes factor greater than 5 indicates that providing sampling dates allowed for successful calibration of the molecular clock, representing strong evidence of a temporal signal (Table 5). The BETS analysis was also conducted on the *CP* and *Rep* genes of both OxYVV and SiYLCV, with (log) Bayes factor values supporting strong evidence of a temporal signal (Table 5).

**Table 5.** Detection of a temporal signal in the begomovirus populations infecting *Sida acuta* in Viçosa, MG, using Bayesian Evaluation of Temporal Signal (BETS). The log marginal likelihood was estimated for both isochronous data (without sampling dates) and heterochronous data (with sampling dates), under both strict and relaxed molecular clock models. The log Bayes factor was calculated as the difference between the log marginal likelihoods of the two competing models. Analyses were conducted for the DNA-A, *Rep*, and *CP* genes of OxYVV and SiYLCV. Log Bayes factors greater than 5 indicate strong statistical support favoring heterochronous models over isochronous ones.

|  | Strict clock | Relaxed clock |
| --- | --- | --- |
| OxYVV DNA-A |  |  |
| Without sampling dates | -9717.87 | -9653.157146 |
| With sampling dates | -9629.50 | -9596.735873 |
| log Bayes factor | 88.37 | 56.42127296 |
| OxYVV *Rep* |  |  |
| Without sampling dates | -3846.46 | -3890.83 |
| With sampling dates | -3777.28 | -3780.82 |
| log Bayes factor | 69.18 | 110.01 |
| OxYVV *CP* |  |  |
| Without sampling dates | -2507.33 | -2509.76 |
| With sampling dates | -2446.65 | -2440.98 |
| log Bayes factor | 60.68 | 68.78 |
| SiYLCV DNA-A |  |  |
| Without sampling dates | -6155.14 | -6148.02 |
| With sampling dates | -6107.14 | -6101.66 |
| log Bayes factor | 47.99 | 46.36 |
| SiYLCV *Rep* |  |  |
| Without sampling dates | -2252.04 | -2266.35 |
| With sampling dates | -2206.64 | -2209.55 |
| log Bayes factor | 45.39 | 56.79 |
| SiYLCV *CP* |  |  |
| Without sampling dates | -1807.04 | -1791.22 |
| With sampling dates | -1754.16 | -1747.22 |
| log Bayes factor | 52.87 | 44.01 |

### Substitution rates of OxYVV and SiYLCV

We estimated the substitution rates per site per year for the DNA-A, *Rep* and *CP* genes of both viruses (Figure 4). The mean substitution rates were in the same order of magnitude across the genomic components of both OxYVV and SiYLCV. For OxYVV, the mean substitution rate was 4.81 × 10⁻⁴ for the DNA-A, 8.26 × 10⁻⁴ for the *CP* gene, and 2.54 × 10⁻⁴ for the *Rep* gene (Figure 4). For SiYLCV, the mean substitution rate was 4.33 × 10⁻⁴ for the DNA-A, 3.17 × 10⁻⁴ for *Rep*, and 1.21 × 10⁻³ for the *CP* gene, the higher value observed (Figure 4).

**Figure 4.**
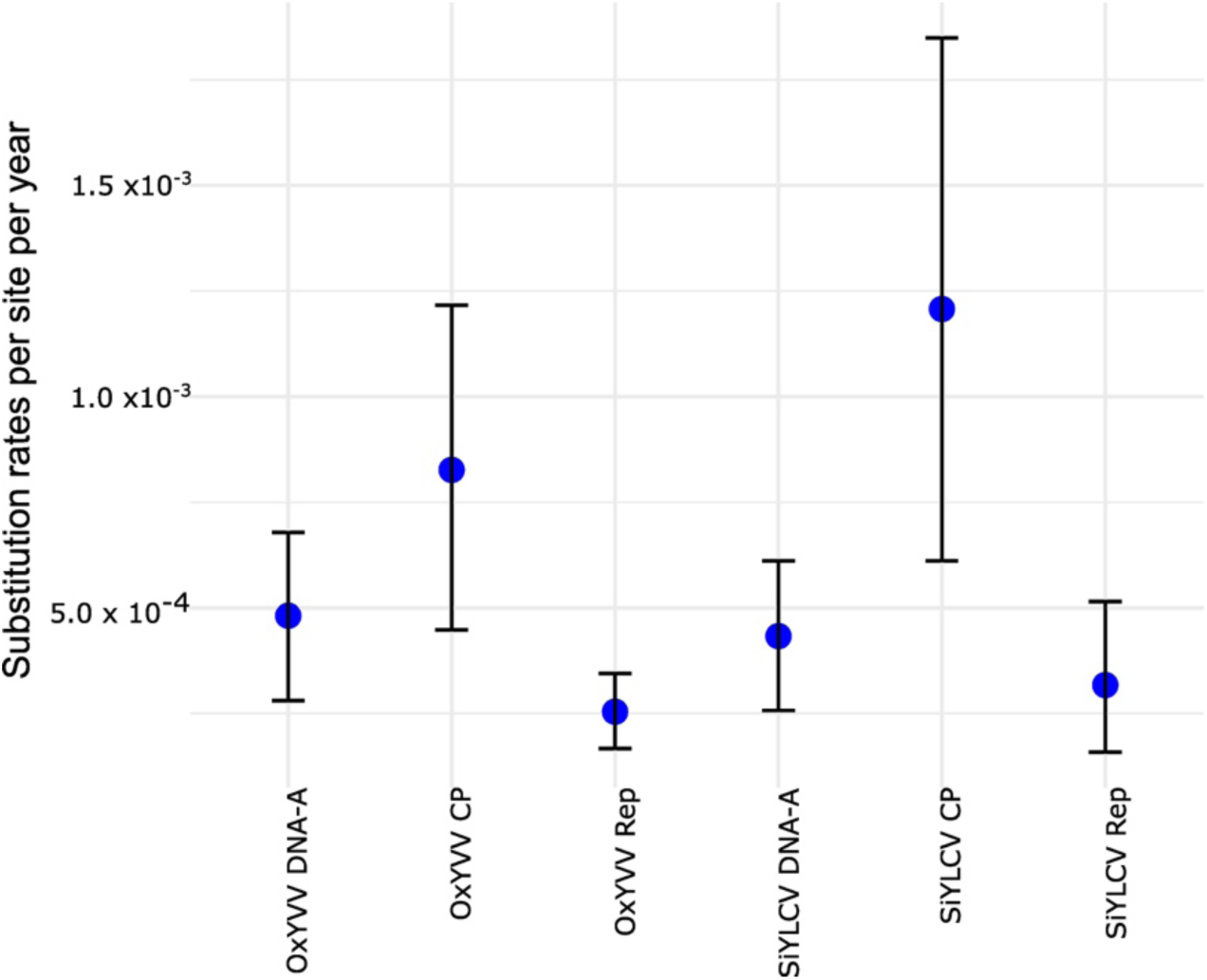
Substitution rates (nucleotides per site per year) for Oxalis yellow vein virus (OxYVV) and Sida yellow leaf curl virus (SiYLCV). Estimated mean substitution rates and corresponding 95% highest posterior density (HPD) intervals are shown for the complete DNA-A, the coat protein (*CP*) gene, and the replication-associated protein (*Rep*) gene of OxYVV and SiYLCV. Estimates were derived using a Bayesian molecular clock framework.

The substitution rates of the complete genomes and the *Rep* genes of the two viruses did not differ statistically (Figure 4). However, for both viruses, the substitution rates of the *CP* gene were significantly higher compared to those of the *Rep* gene. The estimated substitution rates align with those estimated for other begomoviruses (Duffy and Holmes, 2008; Duffy and Holmes, 2009; Jenkins et al., 2002), and are comparable to those observed for RNA viruses.

### Time of the Most Recent Common Ancestor (TMRCA)

The estimated time of the most recent common ancestor (TMRCA) varied across the genomic components of both viruses. For OxYVV, the TMRCA of the complete DNA-A was estimated to be approximately 1931, with a 95% HPD interval between 1858 and 1991 (Figure 5A). The OxYVV *CP* gene had a more recent TMRCA of 1985, with an HPD interval of 1937 to 2005 (Suppl. Figure S7A). The OxYVV *Rep* gene showed the oldest TMRCA, estimated at 1825, with an HPD interval of 1730 to 1888 (Suppl. Figure S8A). For SiYLCV, the TMRCA of the complete DNA-A component was estimated at 1991, with a 95% HPD interval ranging from 1965 to 2005 (Figure 5B). The SiYLCV *CP* gene had the most recent TMRCA, estimated at 2008, with an HPD interval of 1990 to 2011 (Suppl. Figure S7B). The SiYLCV *Rep* gene TMRCA was estimated to be 1995, with a 95% HPD interval between 1975 and 2007 (Suppl. Figure S8B).

**Figure 5.**
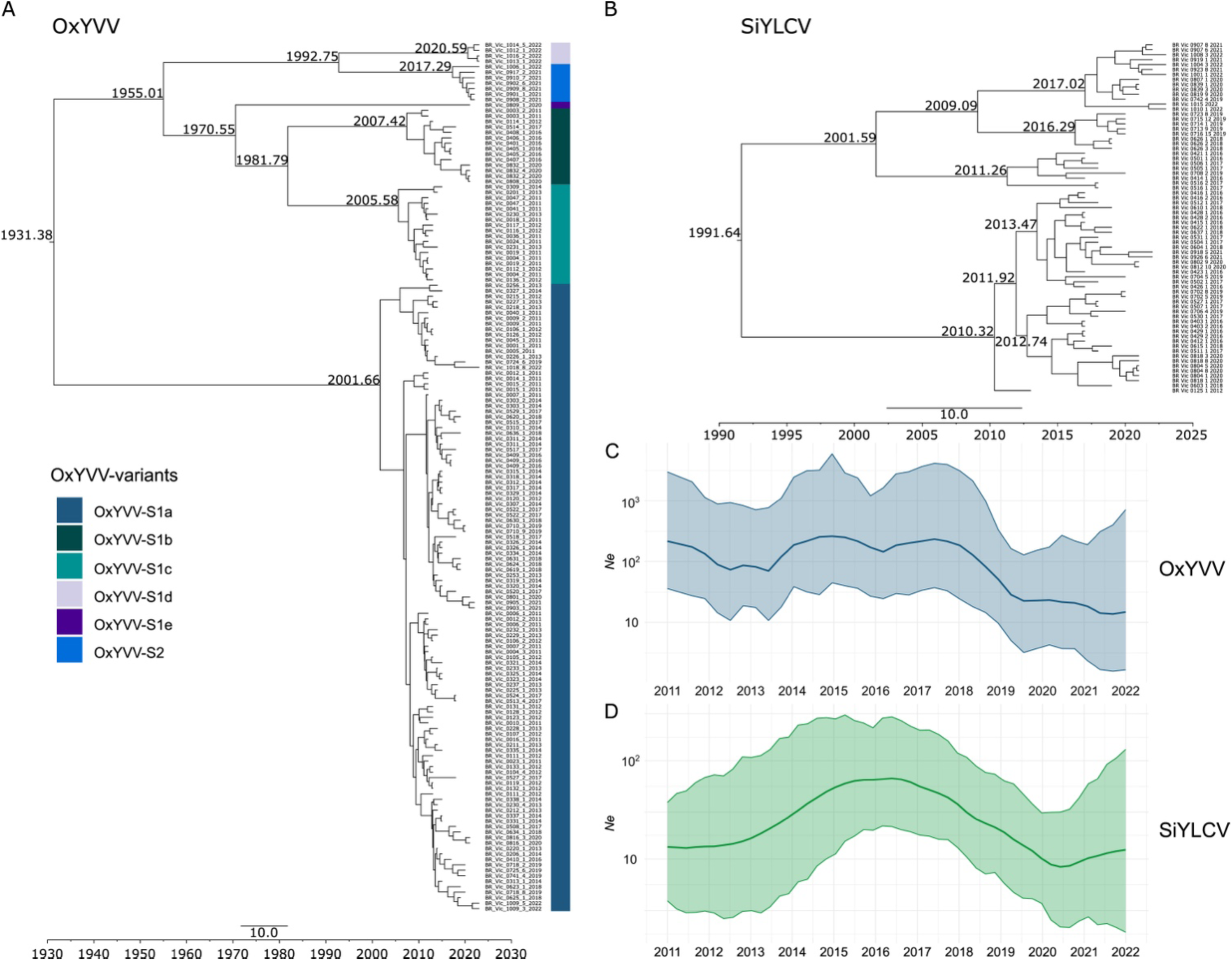
Evolutionary and demographic dynamics of Oxalis yellow vein virus (OxYVV) and Sida yellow leaf curl virus (SiYLCV). **A.** Maximum clade credibility (MCC) phylogenetic tree for OxYVV. The numbers on the branches indicate the time to the most recent common ancestor. The identified OxYVV variants (S1a–S1e and S2) are shown in distinct colors in the side bar. **B.** MCC tree for SiYLCV. **C, D.** Demographic history under a coalescent framework (effective population size, Ne) inferred using the Bayesian Skygrid model for OxYVV (**C**) and SiYLCV (**D**). The lines represent the median estimates, and the shaded area corresponds to the 95% highest posterior density (HPD) intervals.

### Demographic history in a coalescent framework

We hypothesized that the shifts in the composition of the viral community could have been caused by an event that led to a reduction in the effective population size of OxYVV, the prevalent species in the first four years. This may have created an opportunity for SiYLCV to outcompete OxYVV. To test this hypothesis, we reconstructed the demographic trajectory of OxYVV and SiYLCV populations through time using the Bayesian Skygrid Coalescent Model (Hill and Baele, 2019).

The OxYVV population exhibited expansion and contraction dynamics. The first expansion occurred around 2010, reaching a peak in 2016, followed by a decline that extended from mid-2017 to 2022 (Figure 5C). However, there is a degree of uncertainty in the estimation of Ne, which limits our ability to draw conclusions about possible bottlenecks that could explain the changes observed in 2016. We also measured changes in effective population size for the OxYVV *Rep* and OxYVV *CP* genes. The OxYVV *Rep* gene showed a pattern similar to that observed for the complete genome (Suppl. Figure S8C). In contrast, the OxYVV *CP* gene exhibited an almost constant effective population size over time (Suppl. Figure S7C).

SiYLCV showed an increase in effective population size from 2012 to 2016, followed by a reduction after 2016 (Figure 5D). The pattern for the SiYLCV *Rep* gene resembled that of the DNA-A (Suppl. Figure S8C), whereas the *CP* gene showed a nearly constant effective population size over time (Suppl. Figure S7C). The population dynamics captured by the Skygrid model reflects the variation in the genetic diversity of OxYVV and SiYLCV populations over time, and is consistent with the hypothesis that a decline in the OxYVV population allowed for the increase in SiYLCV.

### Evidence of purifying selection and one gene under diversifying selection

Neutrality tests were performed to investigate the role of selection in different coding regions. The tests were performed for the OxYVV and SiYLCV data sets but not for SimVV, due to the small number of sequences in this data set. We obtained negative and significant values for Tajima’s D, Fu & Li D*, and Fu & Li F* tests in most genes of OxYVV and SiYLCV (Suppl. Table S3), indicating an abundance of rare alleles in the populations and suggesting that these genes underwent purifying selection or a recent expansion, as observed in the previous section with the Skygrid analysis.

To identify the type of selection that is acting at the amino acid level in each region of the genome, we estimated ω (dN/dS) using the SLAC method. Most genes showed values of ω <1, indicating that purifying selection is acting on them (Suppl. Table S3). The only exception was the *AC4* gene, which is under diversifying selection for both OxYVV and SiYLCV (Suppl. Table S3). Furthermore, we observed that the intensity of selection on each of the genes varied. The OxYVV *CP*, *Rep*, *MP* and *NSP* genes are under strong negative selection, indicated by very low values of ω. On the other hand, the OxYVV *TrAP* and *REn* genes seem to experience more relaxed negative selection (Suppl. Table S3). The SiYLCV *CP*, *MP* and *NSP* genes are also subject to strong purifying selection (Suppl. Table S3).

Using SLAC, we identified several amino acid sites under negative selection and none under positive selection in the data sets. Using FUBAR, we were able to identify a larger number of sites under selection, including those detected by SLAC and several others also under negative selection, in addition to detecting some sites under positive selection (Suppl. Table S3). We also found several sites under positive selection using MEME. In general, the *CP*, *Rep* and *MP* genes exhibited a higher number of sites under selection, with OxYVV presenting the largest number of sites (Suppl. Table S3). Together, these results indicate that the two viruses experience slightly different selection regimes, evidenced by the number of sites detected in each of the data sets.

## Discussion

Systematic sampling and sequencing of an organism over time provides information that allows the tracking of changes in allele frequencies over generations, the identification of marks of natural selection, and the detection of lateral transfer of genomic regions, giving us an overview of the action of evolutionary mechanisms and their impacts on populations (Billard et al., 2023; Bondaryuk et al., 2023; Ghafari et al., 2024; Mühlemann et al., 2018; Preska Steinberg et al., 2023). Over more than a decade (2011-2022) we monitored a community of begomoviruses infecting *Sida acuta*, a non-cultivated host, in a small area with little, if any, human intervention. Remarkably, even in such a small area we were able to observe a complex dynamics that involved several changes in terms of species, strain and variant composition within the landscape. And it could even be argued that our experimental approach, based on cloning and Sanger sequencing, probably underestimated the extant diversity in the area. However, this approach was useful, or even required, considering the limitations of short-read sequencing in the assembly of complete genomes from reads with >98% identity (which is the level of identity amng the five OxYVV variants) (Golyaev et al., 2025).

The importance of plants from the *Malvaceae* family as begomovirus hosts has been consistently reinforced (Costa, 1955; Ferro et al., 2017a; Fiallo-Olivé et al., 2010; Lima et al., 2021; Macedo et al., 2020; Passos et al., 2017; Pinto et al., 2016). Our results suggest that begomoviruses in *Sida acuta* co-exist in the form of a complex community formed by populations of three viruses (OxYVV, SiYLCV and SimMV) subdivided into strains and variants, plus at least two more viruses (SiMV and MaYVV) present at a lower frequency. Thus, *Sida acuta* is a “mixing vessel” host, which are important players from an evolutionary and ecological perspective, serving as reservoirs and sources of viral genetic diversity (García-Arenal and Zerbini, 2019).

SimMV showed the highest genetic variability, followed by OxYVV, while SiYLCV exhibited very low variability. These patterns were consistent across the two genomic components, DNA-A and DNA-B. Moreover, the DNA-B components of all three viruses showed a higher degree of variability than their cognate DNA-A components, as previously reported for other begomoviruses infecting both non-cultivated and cultivated hosts (Briddon et al., 2010; Xavier et al., 2021). There is direct and indirect evidence pointing to a differential accumulation between the two components, favoring the DNA-B (Pinto et al., 2021; Xiao et al., 2023). The higher the replication rate, the greater the chance of incorrect incorporation of bases (Duffy and Holmes, 2008). In addition, differential accumulation of the two components could result in a reduction of the bottleneck effect for the DNA-B (Billard et al., 2023; Pinto et al., 2021). Lastly, a large region of the DNA-A contains overlapping genes, which could impose strong selection pressure against mutations in these regions (Chen et al., 2019; Kutnjak et al., 2017; Xiao et al., 2023). It is remarkable that, despite infecting the same host in the same area, these viruses presented such distinct variability. These results are suggestive of distinct evolutionary trajectories for the three viruses. In this regard, previous work demonstrated that the genetic variability of begomoviruses is independent of the nature of the host (Mar et al., 2017; Ramos-Sobrinho et al., 2014; Rocha et al., 2013).

Recombination is another important mechanism that generates genetic variability in begomoviruses, contributing to their rapid adaptation to new hosts and environmental conditions (Belabess et al., 2015; Crespo-Bellido et al., 2021; Davino et al., 2009; Lefeuvre and Moriones, 2015). We identified highly reliable recombination events for both the DNA-A and DNA-B in all viruses, albeit at different proportions. Interestingly, two recombination events, one in the DNA-A and the other in the DNA-B and both with OxYVV as the minor parent, were identified in all SiYLCV isolates. Recombination has been shown to be involved in the emergence of well adapted begomoviruses (Belabess et al., 2015; Davino et al., 2009; Monci et al., 2002; Padidam et al., 1999; Pita et al., 2001). It is not unreasonable to propose that a recombination event between a SiYLCV-like parental and OxYVV generated the local SiYLCV population, and that this population has increased fitness compared with both parentals, allowing its co-existence with, and eventually its prevalence over, the local OxYVV population. Further studies assessing the relative fitness of OxYVV and SiYLCV isolates in *Sida acuta* will be conducted to test this hypothesis.

Three recombination events were identified in the SimMV DNA-A, revealing the highly recombinant nature of this species. SimMV has a wide host range, which increases the probability of mixed infections, a prerequisite for recombination (Fernandes et al., 2009; Fontenele et al., 2018; Jovel et al., 2004).

The phylogenetic network constructed for OxYVV showed little reticulation, and indeed a single recombinant individual was identified in the OxYVV population. To date, OxYVV has been reported only from *Oxalis debilis* (Herrera et al., 2015) and *Sida acuta* (this work), and the breadth of its host range is not well known. Nevertheless, *Sida acuta* is a host of several begomoviruses, which favors recombination events due to frequent mixed infections. Added to the high rates of recombination normally detected in begomoviruses (Padidam et al., 1999; Rocha et al., 2013), the fact that only a single recombinant sequence of OxYVV was found in our samples is surprising. It is possible that intraspecific recombination occurs between highly genetically similar individuals (>97% identity), which would make detection by the alignment-based methods in RDP very difficult. Alternatively, a virus that is extremely well adapted to its host may not generate recombinants with improved fitness (or do so at an exceedingly low frequency), and thus the recombinants which are generated, with lower fitness, will be purged from the population. In support of this hypothesis, no intraspecific recombination events were detected in populations of African cassava mosaic virus, bean golden mosaic virus and tomato severe rugose virus, three begomoviruses which are very well adapted to their hosts (cassava, common bean and tomato, respectively) (Lima et al., 2017; Xavier et al., 2021). However, how well adapted OxYVV is to Sida acuta is still not fully understood. Although it is tempting to assume so, given the high frequency with which we found it associated with this host, our sampling area was limited and may not reflect a broader pattern.

Neutrality tests deviated from the hypothesis of neutral selection acting on populations, indicating evidence of negative selection. The ω estimates point to strong evidence of purifying selection in most genes, except for *AC4*, which has been shown to be under diversifying selection in many begomoviruses (Deom et al., 2021; Medina-Puche et al., 2021). *AC4* overlaps in its entirety with *Rep*. Overlapping genes can experience different types of selection, and usually the gene that is involved in accessory functions evolves freely, while the other involved in essential functions experiences more rigorous selective restrictions in order to preserve protein function (Deom et al., 2021; Martín-Hernández and Pagán, 2022; Rancurel et al., 2009). The AC4 protein has distinct functions in the viral infection cycle, which can vary depending on the virus (Medina-Puche et al., 2021). It was shown to be a suppressor of different host defense pathways (Li et al., 2019; Mei et al., 2020; Mei et al., 2021; Vanitharani et al., 2004), but also to confer drought tolerance to some hosts (Corrales-Gutierrez et al., 2020; Medina-Puche et al., 2020). It would be interesting to determine the precise function(s) of AC4 in the infection of *S. acuta* by OxYVV, which could shed some light on the frequent association of begomoviruses with species of the *Sida* genus. Interestingly, amino acid sites under positive selection were identified in all OxYVV genes using MEME, and some were identified in three SiYLCV genes.

This is the first study to evaluate the evolutionary dynamics of begomoviruses in a context that is relatively free from anthropogenic activity, with systematic sampling conducted at the same location and during the same time of year over multiple years. Nevertheless, the estimated substitution rates for OxYVV and SiYLCV were similar to those previously reported for other begomoviruses (Duffy and Holmes, 2008; Duffy and Holmes, 2009), indicating that the evolutionary rate of these viruses in natural environments does not differ significantly from that observed in crop systems. This suggests that such rates may, at least in part, reflect intrinsic evolutionary characteristics of the virus. Furthermore, as the study was conducted in a geographically restricted area yet yielded substitution rates comparable to those observed in widely distributed populations, it also weakens the criticism that previously reported high rates could have been inflated due to artifacts arising from population structure across geographically distinct sampling sites (Allen et al., 2015).

Perhaps the most intriguing observation from our study is the increase in species diversity in the area, from a single virus in 2011 to five viruses in 2021, with a corresponding doubling of the KHILL value during this period. The number of species remained constant from 2011 until 2014, with a sudden change in 2016, followed by a steady increase until 2021. A hypothesis to explain the changes in species composition observed in 2016 would be the occurrence of a random event which resulted in a drastic reduction in the effective size of the OxYVV population. This genetic bottleneck could have altered the adaptive landscape, creating an opportunity for SiYLCV to establish itself in the area (Gallet et al., 2018; Zwart and Elena, 2015). However, the variations in effective population size over time do not provide any evidence of a genetic bottleneck that would have significantly reduced the diversity of OxYVV. On the other hand, we observed the aforementioned recombination event in all SiYLCV isolates, which could explain the shift in viral composition. In this scenario, OxYVV would be the best adapted (and thus prevalent) virus until 2014, when the recombinant SiYLCV emerged and outnumbered OxYVV, the assumption being that SiYLCV is better adapted to *S. acuta* than OxYVV (something that has not been demonstrated experimentally). Moreover, this scenario does not provide an adequate explanation for the further emergence of SimMV, SiMV and MaYVV. A third hypothesis would be that changes in the local whitefly population could have favored the transmission of SiYLCV in relation to OxYVV. It is known that the efficiency of begomovirus transmission can vary among the species of the *Bemisia tabaci* complex (Gautam et al., 2022; Gottlieb et al., 2010). However, we always noticed a very small number of whiteflies in the area, something that did not change during the study (and thus argues against a major change in whitefly populations). But we did not sample whiteflies, and therefore do not have information about the temporal dynamics of the vector in the area. Finally, we should not disregard the possibility of gene flow, which would be a reasonable explanation for the emergence of SimMV, SiMV and MaYVV. Expanding the sampling area would be of value as it could identify source populations. Another alternative would be to explore the dynamics of the landscape, sampling other hosts present in the location - perhaps these sources are not so far away. Possible differences in fitness among the viruses and variants deserve to be biologically tested. Competition assays between the viruses, strains and variants in different contexts can help answer many of the questions that are still open. Furthermore, construction of infectious clones of SiYLCV containing the reversal of the recombination event could clarify whether this event actually culminated in the displacement of OxYVV. These are the first contributions of a major effort to understand ecological and evolutionary processes of begomovirus communities infecting wild hosts outside the agricultural context.

## Acknowledgements

This work was funded by CAPES (Finance Code 001), CNPq (grants 409599/2016-6 and 408159/2023-5 to FMZ) and FAPEMIG (grants APQ-03444-16 and APQ-03278-18 to FMZ). The authors wish to thank Eduardo S.G. Mizubuti and Poliane Alfenas-Zerbini for helpful discussions.

## Notes

### Competing Interest Statement

The authors have declared no competing interest.

## References

Alberdi, A., & Gilbert, M. T. P. (2019). A guide to the application of Hill numbers to DNA-based diversity analyses. Molecular Ecology Resources, 19, 804–817.

Allen, B., Sample, C., Dementieva, Y., Medeiros, R. C., Paoletti, C., & Nowak, M. A. (2015). The molecular clock of neutral evolution can be accelerated or slowed by asymmetric spatial structure. PLoS Computational Biology, 11, e1004108.

Altschul, S. F., Gish, W., Miller, W., Myers, E. W., & Lipman, D. J. (1990). Basic local alignment search tool. Journal of Molecular Biology, 215, 403–410.

Baele, G., Lemey, P., & Suchard, M. A. (2015). Genealogical working distributions for Bayesian model testing with phylogenetic uncertainty. Systematic Biology, 65, 250–264.

Belabess, Z., Dallot, S., El-Montaser, S., Granier, M., Majde, M., Tahiri, A., … Peterschmitt, M. (2015). Monitoring the dynamics of emergence of a non-canonical recombinant of tomato yellow leaf curl virus and displacement of its parental viruses in tomato. Virology, 486, 291–306.

Belabess, Z., Peterschmitt, M., Granier, M., Tahiri, A., Blenzar, A., & Urbino, C. (2016). The non-canonical tomato yellow leaf curl virus recombinant that displaced its parental viruses in Southern Morocco exhibits a high selective advantage in experimental conditions. Journal of General Virology, 97, 3433–3445.

Billard, E., Barro, M., Sérémé, D., Bangratz, M., Wonni, I., Koala, M., … Tollenaere, C. (2023). Dynamics of the rice yellow mottle disease in western Burkina Faso: Epidemic monitoring, spatio-temporal variation of viral diversity, and pathogenicity in a disease hotspot. Virus Evolution, 9, vead049.

Bondaryuk, A. N., Belykh, O. I., Andaev, E. I., & Bukin, Y. S. (2023). Inferring evolutionary timescale of Omsk hemorrhagic fever virus. Viruses, 15, 1576.

Briddon, R. W., Patil, B. L., Bagewadi, B., Nawaz-ul-Rehman, M. S., & Fauquet, C. M. (2010). Distinct evolutionary histories of the DNA-A and DNA-B components of bipartite begomoviruses. BMC Evolutionary Biology, 10, 97.

Brown, J. K., Zerbini, F. M., Navas-Castillo, J., Moriones, E., Ramos-Sobrinho, R., Silva, J. C., … Varsani, A. (2015). Revision of *Begomovirus* taxonomy based on pairwise sequence comparisons. Archives of Virology, 160, 1593–1619.

Campbell, L. I., Nwezeobi, J., van Brunschot, S. L., Kaweesi, T., Seal, S. E., Swamy, R. A. R., … Colvin, J. (2023). Comparative evolutionary analyses of eight whitefly *Bemisia tabac*i sensu lato genomes: Cryptic species, agricultural pests and plant-virus vectors. BMC Genomics, 24, 408.

Castillo-Urquiza, G. P., Beserra Jr., J. E. A., Bruckner, F. P., Lima, A. T. M., Varsani, A., Alfenas-Zerbini, P., & Zerbini, F. M. (2008). Six novel begomoviruses infecting tomato and associated weeds in Southeastern Brazil. Archives of Virology, 153, 1985–1989.

Caulfield, J. L., Wishnok, J. S., & Tannenbaum, S. R. (1998). Nitric oxide-induced deamination of cytosine and guanine in deoxynucleosides and oligonucleotides. Journal of Biological Chemistry, 273, 12689–12695.

Chen, K., Khatabi, B., & Fondong, V. N. (2019). The AC4 protein of a cassava geminivirus is required for virus infection. Molecular Plant-Microbe Interactions, 32, 865–875.

Corrales-Gutierrez, M., Medina-Puche, L., Yu, Y., Wang, L., Ding, X., Luna, A. P., … Lozano-Duran, R. (2020). The C4 protein from the geminivirus *Tomato yellow leaf curl virus* confers drought tolerance in Arabidopsis through an ABA-independent mechanism. Plant Biotechnology Journal, 18, 1121–1123.

Costa, A. S. (1955). Studies on *Abutilon* mosaic in Brazil. Phytopathologische Zeitschrift, 24, 97–112.

Crespo-Bellido, A., Hoyer, J. S., Dubey, D., Jeannot, R. B., & Duffy, S. (2021). Interspecies recombination has driven the macroevolution of cassava mosaic begomoviruses. Journal of Virology, 95, e00541–00521.

Darriba, D., Posada, D., Kozlov, A. M., Stamatakis, A., Morel, B., & Flouri, T. (2020). ModelTest-NG: A new and scalable tool for the selection of DNA and protein evolutionary models. Molecular Biology and Evolution, 37, 291–294.

Davino, S., Napoli, C., Dellacroce, C., Miozzi, L., Noris, E., Davino, M., & Accotto, G. P. (2009). Two new natural begomovirus recombinants associated with the tomato yellow leaf curl disease co-exist with parental viruses in tomato epidemics in Italy. Virus Research, 143, 15–23.

De Barro, P. J., Liu, S. S., Boykin, L. M., & Dinsdale, A. B. (2011). *Bemisia tabaci*: A statement of species status. Annual Review of Entomology, 56, 1–19.

Deom, C. M., Brewer, M. T., & Severns, P. M. (2021). Positive selection and intrinsic disorder are associated with multifunctional C4(AC4) proteins and geminivirus diversification. Scientific Reports, 11, 11150.

Dolan, P. T., Whitfield, Z. J., & Andino, R. (2018). Mechanisms and concepts in RNA virus population dynamics and evolution. Annual Review of Virology, 5, 69–92.

Doyle, J. J., & Doyle, J. L. (1987). A rapid DNA isolation procedure for small amounts of fresh leaf tissue. Phytochemical Bulletin, 19, 11–15.

Drummond, A. J., & Rambaut, A. (2007). BEAST: Bayesian evolutionary analysis by sampling trees. BMC Evolutionary Biology, 7, 214.

Duchene, S., Lemey, P., Stadler, T., Ho, S. Y. W., Duchene, D. A., Dhanasekaran, V., & Baele, G. (2020). Bayesian Evaluation of Temporal Signal in measurably evolving populations. Molecular Biology and Evolution, 37, 3363–3379.

Duffy, S., & Holmes, E. C. (2008). Phylogenetic evidence for rapid rates of molecular evolution in the single-stranded DNA begomovirus tomato yellow leaf curl virus. Journal of Virology, 82, 957–965.

Duffy, S., & Holmes, E. C. (2009). Validation of high rates of nucleotide substitution in geminiviruses: Phylogenetic evidence from East African cassava mosaic viruses. Journal of General Virology, 90, 1539–1547.

Duffy, S., Shackelton, L. A., & Holmes, E. C. (2008). Rates of evolutionary change in viruses: Patterns and determinants. Nature Reviews Genetics, 9, 267–276.

Fernandes, F. R., Albuquerque, L. C., Giordano, L. B., Boiteux, L. S., Ávila, A. C., & Inoue-Nagata, A. K. (2008). Diversity and prevalence of Brazilian bipartite begomovirus species associated to tomatoes. Virus Genes, 36, 251–258.

Fernandes, F. R., Cruz, A. R. R., Faria, J. C., Zerbini, F. M., & Aragão, F. J. L. (2009). Three distinct begomoviruses associated with soybean in central Brazil. Archives of Virology, 154, 1567–1570.

Ferro, C. G., Silva, J. P., Xavier, C. A. D., Godinho, M. T., Lima, A. T. M., Mar, T. B., … Zerbini, F. M. (2017a). The ever increasing diversity of begomoviruses infecting non-cultivated hosts: new species from *Sida* spp. and *Leonurus sibiricus*, plus two New World alphasatellites. Annals of Applied Biology, 170, 204–218.

Ferro, M. M. M., Ramos-Sobrinho, R., Silva, J. T., Assunção, I. P., & Lima, G. S. A. (2017b). Genetic structure of populations of the begomoviruses *Tomato mottle leaf curl virus* and *Sida mottle Alagoas virus* infecting tomato (*Solanum lycopersicum*) and *Sida* spp., respectively. Tropical Plant Pathology, 42, 39–45.

Fiallo-Olive, E., Lett, J. M., Martin, D. P., Roumagnac, P., Varsani, A., Zerbini, F. M., & Navas-Castillo, J. (2021). ICTV Virus Taxonomy Profile: *Geminiviridae* 2021. Journal of General Virology, 102, 001696.

Fiallo-Olivé, E., Martinez-Zubiaur, Y., Moriones, E., & Navas-Castillo, J. (2010). Complete nucleotide sequence of *Sida golden mosaic Florida virus* and phylogenetic relationships with other begomoviruses infecting malvaceous weeds in the Caribbean. Archives of Virology, 155, 1535–1537.

Fiallo-Olivé, E., Trenado, H. P., Louro, D., & Navas-Castillo, J. (2019). Recurrent speciation of a tomato yellow leaf curl geminivirus in Portugal by recombination. Scientific Reports, 9, 1–8.

Fontenele, R. S., Ribeiro, G. C., Lamas, N. S., Ribeiro, S. G., Costa, A. F., Boiteux, L. S., & Fonseca, M. E. N. (2018). First report of *Sida micrantha mosaic virus* Infecting *Oxalis* species in Brazil. Plant Disease, 102, 1862.

Fu, L., Niu, B., Zhu, Z., Wu, S., & Li, W. (2012). CD-HIT: accelerated for clustering the next-generation sequencing data. Bioinformatics, 28, 3150–3152.

Gallet, R., Fabre, F., Thebaud, G., Sofonea, M. T., Sicard, A., Blanc, S., & Michalakis, Y. (2018). Small bottleneck size in a highly multipartite virus during a complete infection cycle. Journal of Virology, 92, e00139–00118.

García-Arenal, F., & Zerbini, F. M. (2019). Life on the edge: Geminiviruses at the interface between crops and wild plant hosts. Annual Review of Virology, 6, 411–433.

Gautam, S., Mugerwa, H., Buck, J. W., Dutta, B., Coolong, T., Adkins, S., & Srinivasan, R. (2022). Differential transmission of Old and New World begomoviruses by Middle East-Asia Minor 1 (MEAM1) and Mediterranean (MED) cryptic species of *Bemisia tabaci*. Viruses, 14, 1104.

Ghafari, M., Sõmera, M., Sarmiento, C., Niehl, A., Hébrard, E., Tsoleridis, T., … Fargette, D. (2024). Revisiting the origins of the *Sobemovirus* genus: A case for ancient origins of plant viruses. PLoS Pathogens, 20, e1011911.

Golyaev, V., Dierickx, S., Deforche, K., Dumon, W., & Vanderschuren, H. (2025). A method for in-depth analysis of circular DNA virus populations by unambiguously profiling the low abundant virus variants and partial genomic components. Nucleic Acids Research, 53.

Gottlieb, Y., Zchori-Fein, E., Mozes-Daube, N., Kontsedalov, S., Skaljac, M., Brumin, M., … Ghanim, M. (2010). The transmission efficiency of tomato yellow leaf curl virus by the whitefly *Bemisia tabaci* is correlated with the presence of a specific symbiotic bacterium species. Journal of Virology, 84, 9310–9317.

Hahm, J. Y., Park, J., Jang, E. S., & Chi, S. W. (2022). 8-Oxoguanine: from oxidative damage to epigenetic and epitranscriptional modification. Experimental and Molecular Medicine, 54, 1626–1642.

Hanley-Bowdoin, L., Bejarano, E. R., Robertson, D., & Mansoor, S. (2013). Geminiviruses: Masters at redirecting and reprogramming plant processes. Nature Reviews Microbiology, 11, 777–788.

Herrera, F., Aboughanem-Sabanadzovic, N., & Valverde, R. A. (2015). A begomovirus associated with yellow vein symptoms of *Oxalis debilis*. European Journal of Plant Pathology, 142, 203–208.

Hesketh, E. L., Saunders, K., Fisher, C., Potze, J., Stanley, J., Lomonossoff, G. P., & Ranson, N. A. (2018). The 3.3 Å structure of a plant geminivirus using cryo-EM. Nature Communications, 9, 2369.

Hill, M. O. (1973). Diversity and evenness - Unifying notation and its consequences. Ecology, 54, 427–432.

Hill, V., & Baele, G. (2019). Bayesian estimation of past population dynamics in BEAST 1.10 using the Skygrid coalescent model. Molecular Biology and Evolution, 36, 2620–2628.

Hsieh, T. C., Ma, K. H., & Chao, A. (2016). iNEXT: An R package for rarefaction and extrapolation of species diversity (Hill numbers). Methods in Ecology and Evolution, 7, 1451–1456.

Huson, D. H., & Bryant, D. (2006). Application of phylogenetic networks in evolutionary studies. Molecular Biology and Evolution, 23, 254–267.

Inoue-Nagata, A. K., Albuquerque, L. C., Rocha, W. B., & Nagata, T. (2004). A simple method for cloning the complete begomovirus genome using the bacteriophage phi29 DNA polymerase. Journal of Virological Methods, 116, 209–211.

Jenkins, G. M., Rambaut, A., Pybus, O. G., & Holmes, E. C. (2002). Rates of molecular evolution in RNA viruses: A quantitative phylogenetic analysis. Journal of Molecular Evolution, 54, 156–165.

Jost, L. (2006). Entropy and diversity. Oikos, 113, 363–375.

Jovel, J., Reski, G., Rothenstein, D., Ringel, M., Frischmuth, T., & Jeske, H. (2004). *Sida micrantha* mosaic is associated with a complex infection of begomoviruses different from *Abutilon mosaic virus*. Archives of Virology, 149, 829–841.

Kass, R. E., & Raftery, A. E. (1995). Bayes Factors. Journal of the American Statistical Association, 90, 773–795.

Katoh, K., & Standley, D. M. (2013). MAFFT Multiple Sequence Alignment Software Version 7: Improvements in Performance and Usability. Molecular Biology and Evolution, 30, 772–780.

Kearse, M., Moir, R., Wilson, A., Stones-Havas, S., Cheung, M., Sturrock, S., … Drummond, A. (2012). Geneious Basic: An integrated and extendable desktop software platform for the organization and analysis of sequence data. Bioinformatics, 28, 1647–1649.

Kilpatrick, A. M., & Randolph, S. E. (2012). Drivers, dynamics, and control of emerging vector-borne zoonotic diseases. Lancet, 380, 1946–1955.

Kosakovsky-Pond, S. L., & Frost, S. D. W. (2005). Not so different after all: A comparison of methods for detecting amino acid sites under selection. Molecular Biology and Evolution, 22, 1208–1222.

Kozlov, A. M., Darriba, D., Flouri, T., Morel, B., & Stamatakis, A. (2019). RAxML-NG: a fast, scalable and user-friendly tool for maximum likelihood phylogenetic inference. Bioinformatics, 35, 4453–4455.

Kutnjak, D., Elena, S. F., & Ravnikar, M. (2017). Time-sampled population sequencing reveals the interplay of selection and genetic drift in experimental evolution of Potato virus Y. Journal of Virology, 91, e00690–00617.

Lefeuvre, P., & Moriones, E. (2015). Recombination as a motor of host switches and virus emergence: Geminiviruses as case studies. Current Opinion in Virology, 10, 14–19.

Li, T., Huang, Y., Xu, Z. S., Wang, F., & Xiong, A. S. (2019). Salicylic acid-induced differential resistance to the Tomato yellow leaf curl virus among resistant and susceptible tomato cultivars. BMC Plant Biology, 19, 173.

Lima, A. T. M., Orílio, A. F., Almeida, M. M. S., Rocha, C. S., Barros, D. R., Castillo-Urquiza, G. P., … Zerbini, F. M. (2021). Malvaviscus yellow mosaic virus, a divergent begomovirus carrying a nanovirus-like nonanucleotide and a modified stem-loop structure. Annals of Applied Biology, 179, 96–107.

Lima, A. T. M., Silva, J. C. F., Silva, F. N., Castillo-Urquiza, G. P., Silva, F. F., Seah, Y. M., … Zerbini, F. M. (2017). The diversification of begomovirus populations is predominantly driven by mutational dynamics. Virus Evolution, 3, vex005.

Macedo, M. A., Rego-Machado, C. M., Maliano, M. L., Rojas, M. R., Inoue-Nagata, A. K., & Gilbertson, R. L. (2020). Complete sequence of a new bipartite begomovirus infecting *Sida* sp. in Northeastern Brazil. Archives of Virology, 165, 253–256.

Mar, T. B., Xavier, C. A. D., Lima, A. T. M., Nogueira, A. M., Silva, J. C. F., Ramos-Sobrinho, R., … Zerbini, F. M. (2017). Genetic variability and population structure of the New World begomovirus *Euphorbia yellow mosaic virus*. Journal of General Virology, 98, 1537–1551.

Martin, D. P., Varsani, A., Roumagnac, P., Botha, G., Maslamoney, S., Schwab, T., … Murrell, B. (2021). RDP5: A computer program for analyzing recombination in, and removing signals of recombination from, nucleotide sequence datasets. Virus Evolution, 7, veaa087.

Martín-Hernández, I., & Pagán, I. (2022). Gene overlapping as a modulator of begomovirus evolution. Microorganisms, 10.

McVean, G., Awadalla, P., & Fearnhead, P. (2002). A coalescent-based method for detecting and estimating recombination from gene sequences. Genetics, 160, 1231–1241.

Medina-Puche, L., Orilio, A. F., Zerbini, F. M., & Lozano-Duran, R. (2021). Small but mighty: Functional landscape of the versatile geminivirus-encoded C4 protein. PLoS Pathogens, 17, e1009915.

Medina-Puche, L., Tan, H., Dogra, V., Wu, M., Rosas-Diaz, T., Wang, L., … Lozano-Duran, R. (2020). A defense pathway linking plasma membrane and chloroplasts and co-opted by pathogens. Cell, 182, 1109–1124.e1125.

Mei, Y., Ma, Z., Wang, Y., & Zhou, X. (2020). Geminivirus C4 antagonizes the HIR1-mediated hypersensitive response by inhibiting the HIR1 self-interaction and promoting degradation of the protein. New Phytologist, 225, 1311–1326.

Mei, Y., Wang, Y., Hu, T., He, Z., & Zhou, X. (2021). The C4 protein encoded by *Tomato leaf curl Yunnan virus* interferes with mitogen-activated protein kinase cascade-related defense responses through inhibiting the dissociation of the ERECTA/BKI1 complex. New Phytologist, 231, 747–762.

Monci, F., Sanchez-Campos, S., Navas-Castillo, J., & Moriones, E. (2002). A natural recombinant between the geminiviruses tomato yellow leaf curl Sardinia virus and tomato yellow leaf curl virus exhibits a novel pathogenic phenotype and is becoming prevalent in Spanish populations. Virology, 303, 317–326.

Mühlemann, B., Jones, T. C., Damgaard, P. d. B., Allentoft, M. E., Shevnina, I., Logvin, A., … Willerslev, E. (2018). Ancient hepatitis B viruses from the Bronze Age to the Medieval period. Nature, 557, 418–423.

Murrell, B., Moola, S., Mabona, A., Weighill, T., Sheward, D., Kosakovsky Pond, S. L., & Scheffler, K. (2013). FUBAR: A fast, unconstrained bayesian approximation for inferring selection. Molecular Biology and Evolution, 30, 1196–1205.

Murrell, B., Wertheim, J. O., Moola, S., Weighill, T., Scheffler, K., & Kosakovsky Pond, S. L. (2012). Detecting individual sites subject to episodic diversifying selection. PLoS Genetics, 8, e1002764.

Narechania, A., Bobo, D., Deitz, K., DeSalle, R., Planet, P. J., & Mathema, B. (2024). Rapid SARS-CoV-2 surveillance using clinical, pooled, or wastewater sequence as a sensor for population change. Genome Research, 34, 1651–1660.

Nei, M. (1987). *Molecular Evolutionary Genetics*. New York: Columbia University Press.

Padidam, M., Sawyer, S., & Fauquet, C. M. (1999). Possible emergence of new geminiviruses by frequent recombination. Virology, 265, 218–224.

Passos, L. S., Teixeira, J. W. M., Teixeira, K. J. M. L., Xavier, C. A. D., Zerbini, F. M., Araújo, A. S. F., & Beserra, J. E. A. (2017). Two new begomoviruses that infect non-cultivated malvaceae in Brazil. Archives of Virology, 162, 1795–1797.

Patil, B. L., & Fauquet, C. M. (2009). Cassava mosaic geminiviruses: actual knowledge and perspectives. Molecular Plant Pathology, 10, 685–701.

Pinto, V. B., Quadros, A. F. F., Godinho, M. T., Silva, J. C., Alfenas-Zerbini, P., & Zerbini, F. M. (2021). Intra-host evolution of the ssDNA virus tomato severe rugose virus (ToSRV). Virus Research, 292, 198234.

Pinto, V. B., Silva, J. P., Fiallo-Olivé, E., Navas-Castillo, J., & Zerbini, F. M. (2016). Novel begomoviruses recovered from *Pavonia* sp. in Brazil. Archives of Virology, 161, 735–739.

Pita, J. S., Fondong, V. N., Sangare, A., Otim-Nape, G. W., Ogwal, S., & Fauquet, C. M. (2001). Recombination, pseudorecombination and synergism of geminiviruses are determinant keys to the epidemic of severe cassava mosaic disease in Uganda. Journal of General Virology, 82, 655–665.

Pond, S. L. K., Posada, D., Gravenor, M. B., Woelk, C. H., & Frost, S. D. W. (2006). GARD: a genetic algorithm for recombination detection. Bioinformatics, 22, 3096–3098.

Preska Steinberg, A., Silander, O. K., & Kussell, E. (2023). Correlated substitutions reveal SARS-like coronaviruses recombine frequently with a diverse set of structured gene pools. Proceedings of the National Academy of Sciences of the United States of America, 120, e2206945119.

Pybus, O. G., Tatem, A. J., & Lemey, P. (2015). Virus evolution and transmission in an ever more connected world. Proceedings of the Royal Society B - Biological Sciences, 282, 20142878.

Rambaut, A., Drummond, A. J., Xie, D., Baele, G., & Suchard, M. A. (2018). Posterior summarization in Bayesian phylogenetics using Tracer 1.7. Systematic Biology, 67, 901–904.

Ramos-Sobrinho, R., Xavier, C. A. D., Pereira, H. M. B., Lima, G. S. A., Assunção, I. P., Mizubuti, E. S. G., … Zerbini, F. M. (2014). Contrasting genetic structure between two begomoviruses infecting the same leguminous hosts. Journal of General Virology, 95, 2540–2552.

Rancurel, C., Khosravi, M., Dunker, A. K., Romero, P. R., & Karlin, D. (2009). Overlapping genes produce proteins with unusual sequence properties and offer insight into de novo protein creation. Journal of Virology, 83, 10719–10736.

Ribeiro, S. G., Ávila, A. C., Bezerra, I. C., Fernandes, J. J., Faria, J. C., Lima, M. F., … Zerbini, F. M. (1998). Widespread occurrence of tomato geminiviruses in Brazil, associated with the new biotype of the whitefly vector. Plant Disease, 82, 830.

Rocha, C. S., Castillo-Urquiza, G. P., Lima, A. T. M., Silva, F. N., Xavier, C. A. D., Hora-Junior, B. T., … Zerbini, F. M. (2013). Brazilian begomovirus populations are highly recombinant, rapidly evolving, and segregated based on geographical location. Journal of Virology, 87, 5784–5799.

Rodelo-Urrego, M., Pagán, I., González-Jara, P., Betancourt, M., Moreno-Letelier, A., Ayllón, M. A., … García-Arenal, F. (2013). Landscape heterogeneity shapes host-parasite interactions and results in apparent plant-virus codivergence. Molecular Ecology, 22, 2325–2340.

Rodríguez-Negrete, E. A., Morales-Aguilar, J. J., Domínguez-Duran, G., Torres-Devora, G., Camacho-Beltrán, E., Leyva-López, N. E., … Méndez-Lozano, J. (2019). High-throughput sequencing reveals differential begomovirus species diversity in non-cultivated plants in Northern-Pacific Mexico. Viruses, 11, 594.

Rojas, M. R., Macedo, M. A., Maliano, M. R., Soto-Aguilar, M., Souza, J. O., Briddon, R. W., … Gilbertson, R. L. (2018). World management of geminiviruses. Annual Review of Phytopathology, 56, 637–677.

Roossinck, M. J., & Garcia-Arenal, F. (2015). Ecosystem simplification, biodiversity loss and plant virus emergence. Current Opinion in Virology, 10, 56–62.

Rozas, J., Ferrer-Mata, A., Sánchez-DelBarrio, J. C., Guirao-Rico, S., Librado, P., Ramos-Onsins, S. E., & Sánchez-Gracia, A. (2017). DnaSP 6: DNA sequence polymorphism analysis of large data sets. Molecular Biology and Evolution, 34, 3299–3302.

Sambrook, J., & Russel, D. (2001). *Molecular Cloning - A Laboratory Manual (3^a^ ed.)*. Cold Spring Harbor, NY: Cold Spring Harbor Laboratory Press.

Sánchez-Campos, S., Diaz, J. A., Monci, F., Bejarano, E. R., Reina, J., Navas-Castillo, J., … Moriones, E. (2002). High genetic stability of the begomovirus tomato yellow leaf curl Sardinia virus in southern Spain over an 8-year period. Phytopathology, 92, 842–849.

Sayers, E. W., Bolton, E. E., Brister, J. R., Canese, K., Chan, J., Comeau, D. C., … Sherry, S. T. (2022). Database resources of the National Center for Biotechnology Information. Nucleic Acids Research, 50, D20–d26.

Shakir, S., Zaidi, S. S.-A., Farooq, M., Amin, I., Scheffler, J., Scheffler, B., … Mansoor, S. (2019). Non-cultivated cotton species (*Gossypium* spp.) act as a reservoir for cotton leaf curl begomoviruses and associated satellites. Plants, 8, 127.

Souza, J. O., Melgarejo, T. A., Vu, S., Nakasu, E. Y. T., Chen, L. F., Rojas, M. R., … Gilbertson, R. L. (2022). How to be a successful monopartite begomovirus in a bipartite-dominated world: Emergence and spread of tomato mottle leaf curl virus in Brazil. Journal of Virology, 96, e0072522.

Suchard, M. A., Lemey, P., Baele, G., Ayres, D. L., Drummond, A. J., & Rambaut, A. (2018). Bayesian phylogenetic and phylodynamic data integration using BEAST 1.10. Virus Evolution, 4.

Swofford, D. L. (2003). PAUP*. Phylogenetic Analysis Using Parsimony (*and Other Methods). Version 4. Sunderland, Massachusetts: Sinauer Associates.

van der Walt, E., Martin, D. P., Varsani, A., Polston, J. E., & Rybicki, E. P. (2008). Experimental observations of rapid Maize streak virus evolution reveal a strand-specific nucleotide substitution bias. Virology Journal, 5, 104.

Vanitharani, R., Chellappan, P., Pita, J. S., & Fauquet, C. M. (2004). Differential roles of AC2 and AC4 of cassava geminiviruses in mediating synergism and suppression of posttranscriptional gene silencing. Journal of Virology, 78, 9487–9498.

Weaver, S., Shank, S. D., Spielman, S. J., Li, M., Muse, S. V., & Kosakovsky Pond, S. L. (2018). Datamonkey 2.0: A modern web application for characterizing selective and other evolutionary processes. Molecular Biology and Evolution, 35, 773–777.

Wood, M. L., Esteve, A., Morningstar, M. L., Kuziemko, G. M., & Essigmann, J. M. (1992). Genetic effects of oxidative DNA damage: Comparative mutagenesis of 7,8-dihydro-8-oxoguanine and 7,8-dihydro-8-oxoadenine in Escherichia coli. Nucleic Acids Research, 20, 6023–6032.

Xavier, C. A. D., Godinho, M. T., Mar, T. B., Ferro, C. G., Sande, O. F. L., Silva, J. C., … Zerbini, F. M. (2021). Evolutionary dynamics of bipartite begomoviruses revealed by complete genome analysis. Molecular Ecology, 15, 3747–3767.

Xiao, Y. X., Li, D., Wu, Y. J., Liu, S. S., & Pan, L. L. (2023). Constant ratio between the genomic components of bipartite begomoviruses during infection and transmission. Virology Journal, 20, 186.

Yang, X. L., Zhou, M. N., Qian, Y. J., Xie, Y., & Zhou, X. P. (2014). Molecular variability and evolution of a natural population of *Tomato yellow leaf curl virus* in Shanghai, China. Journal of Zhejiang University-Science B, 15, 133–142.

Zwart, M. P., & Elena, S. F. (2015). Matters of size: genetic bottlenecks in virus infection and their potential impact on evolution. Annual Review of Virology, 2, 161–179.

