## Supplementary Tables S1-S3 and Supplementary Figures S1-S8 for "Evolution of ssDNA plant viruses in the natural environment - a journey through time"

**Supplementary Table S1.** *Sida acuta* samples collected in Viçosa, MG, from 2011 to 2022, and the corresponding begomovirus clones obtained from each sample. Samples indicated in bold had a mixed infection.

| Sample | Year | Isolate | Genbank acc. number | Comp. | Enzyme | Species- variant |
|---|---|---|---|---|---|---|
| 01 | 2011 | BR_Vic_0001_1_2011 | PX277586 | DNA-A | ClaI | OxYVV-S1a |
| | | BR_Vic_0001_1_2011 | PX277748 | DNA-B | BamHI | OxYVV |
| 03 | 2011 | BR_Vic_0003_1_2011 | PX277587 | DNA-A | ClaI | OxYVV-S1b |
| | | BR_Vic_0003_2_2011 | PX277588 | DNA-A | ClaI | OxYVV-S1b |
| | | BR_Vic_0003_1_2011 | PX277769 | DNA-B | BamHI | OxYVV |
| | | BR_Vic_0003_2_2011 | PX277770 | DNA-B | BamHI | OxYVV |
| | | BR_Vic_0003_3_2011 | PX277774 | DNA-B | BamHI | OxYVV |
| 04 | 2011 | BR_Vic_0004_1_2011 | PX277589 | DNA-A | ClaI | OxYVV-S1c |
| | | BR_Vic_0004_2_2011 | PX277590 | DNA-A | EcoRV | OxYVV-S1c |
| | | BR_Vic_0004_3_2011 | PX277591 | DNA-A | ClaI | OxYVV-S1a |
| 05 | 2011 | BR_Vic_0005_2011 | PX277592 | DNA-A | ClaI | OxYVV-S1a |
| | | BR_Vic_0005_3_2011 | PX277759 | DNA-B | BamHI | OxYVV |
| 06 | 2011 | BR_Vic_0006_1_2011 | PX277593 | DNA-A | ClaI | OxYVV-S1a |
| | | BR_Vic_0006_1_2011 | PX277762 | DNA-A | EcoRV | OxYVV-S1a |
| | | BR_Vic_0006_2_2011 | PX277594 | DNA-B | BamHI | OxYVV |
| 07 | 2011 | BR_Vic_0007_1_2011 | PX277595 | DNA-A | ClaI | OxYVV-S1a |
| | | BR_Vic_0007_2_2011 | PX277596 | DNA-A | ClaI | OxYVV-S1a |
| | | BR_Vic_0007_1_2011 | PX277752 | DNA-B | BamHI | OxYVV |
| | | BR_Vic_0007_2_2011 | PX277753 | DNA-B | BamHI | OxYVV |
| 09 | 2011 | BR_Vic_0009_1_2011 | PX277597 | DNA-A | ClaI | OxYVV-S1a |
| | | BR_Vic_0009_2_2011 | PX277598 | DNA-A | EcoRV | OxYVV-S1a |
| | | BR_Vic_0009_1_2011 | PX277749 | DNA-B | BamHI | OxYVV |
| 10 | 2011 | BR_Vic_0010_1_2011 | PX277599 | DNA-A | ClaI | OxYVV-S1a |
| | | BR_Vic_0010_1_2011 | PX277755 | DNA-B | BamHI | OxYVV |
| 12 | 2011 | BR_Vic_0012_1_2011 | PX277600 | DNA-A | ClaI | OxYVV-S1a |
| | | BR_Vic_0012_2_2011 | PX277601 | DNA-A | ClaI | OxYVV-S1a |
| | | BR_Vic_0012_1_2011 | PX277761 | DNA-B | BamHI | OxYVV |
| 13 | 2011 | BR_Vic_0013_1_2011 | PX277756 | DNA-B | BamHI | OxYVV |
| | | BR_Vic_0013_2_2011 | PX277750 | DNA-B | BamHI | OxYVV |
| 14 | 2011 | BR_Vic_0014_1_2011 | PX277602 | DNA-A | ClaI | OxYVV-S1a |
| | | BR_Vic_0014_2_2011 | PX277760 | DNA-B | BamHI | OxYVV |
| 15 | 2011 | BR_Vic_0015_1_2011 | PX277603 | DNA-A | ClaI | OxYVV-S1a |
| | | BR_Vic_0015_2_2011 | PX277604 | DNA-A | ClaI | OxYVV-S1a |
| | | BR_Vic_0015_1_2011 | PX277751 | DNA-B | BamHI | OxYVV |

| | | | | | | |
|---|---|---|---|---|---|---|
| | | BR_Vic_0015_1_2011 | PX277751 | DNA-B | BamHI | OxYVV |
| 16 | 2011 | BR_Vic_0016_1_2011 | PX277605 | DNA-A | ClaI | OxYVV-S1a |
| | | BR_Vic_0016_2_2011 | PX277786 | DNA-B | ClaI | OxYVV-S1a |
| | | BR_Vic_0016_3_2011 | PX277776 | DNA-B | ClaI | OxYVV-S1a |
| 18 | 2011 | BR_Vic_0018_1_2011 | PX277606 | DNA-A | EcoRV | OxYVV-S1c |
| | | BR_Vic_0018_1_2011 | PX277780 | DNA-B | ClaI | OxYVV-S1c |
| 19 | 2011 | BR_Vic_0019_1_2011 | PX277607 | DNA-A | ClaI | OxYVV-S1c |
| | | BR_Vic_0019_2_2011 | PX277608 | DNA-A | ClaI | OxYVV-S1c |
| | | BR_Vic_0019_1_2011 | PX277782 | DNA-B | BamHI | OxYVV-S1c |
| 20 | 2011 | BR_Vic_0020_1_2011 | PX277775 | DNA-B | ClaI | OxYVV |
| 23 | 2011 | BR_Vic_0023_1_2011 | PX277609 | DNA-A | ClaI | OxYVV-S1a |
| | | BR_Vic_0023_1_2011 | PX277754 | DNA-B | BamHI | OxYVV |
| 24 | 2011 | BR_Vic_0024_1_2011 | PX277610 | DNA-A | ClaI | OxYVV-S1c |
| 27 | 2011 | BR_Vic_0027_1_2011 | PX277771 | DNA-B | ClaI | OxYVV |
| 32 | 2011 | BR_Vic_0032_2_2011 | PX277777 | DNA-B | BamHI | OxYVV |
| 36 | 2011 | BR_Vic_0036_1_2011 | PX277611 | DNA-A | ClaI | OxYVV-S1c |
| 40 | 2011 | BR_Vic_0040_1_2011 | PX277612 | DNA-A | ClaI | OxYVV-S1a |
| | | BR_Vic_0040_1_2011 | PX277793 | DNA-B | ClaI | OxYVV-S1a |
| 41 | 2011 | BR_Vic_0041_1_2011 | PX277613 | DNA-A | ClaI | OxYVV-S1c |
| | | BR_Vic_0041_3_2011 | PX277795 | DNA-B | ClaI | OxYVV-S1c |
| 42 | 2011 | BR_Vic_0042_1_2011 | PX277778 | DNA-B | ClaI | OxYVV |
| | | BR_Vic_0042_2_2011 | PX277787 | DNA-B | ClaI | OxYVV |
| 44 | 2011 | BR_Vic_0044_1_2011 | PX277757 | DNA-B | BamHI | OxYVV |
| | | BR_Vic_0044_3_2011 | PX277758 | DNA-B | BamHI | OxYVV |
| 45 | 2011 | BR_Vic_0045_1_2011 | PX277614 | DNA-A | EcoRV | OxYVV-S1a |
| 46 | 2011 | BR_Vic_0046_1_2011 | PX277779 | DNA-B | BamHI | OxYVV |
| | | BR_Vic_0046_2_2011 | PX277781 | DNA-B | ClaI | OxYVV |
| 47 | 2011 | BR_Vic_0047_1_2011 | PX277615 | DNA-A | ClaI | OxYVV-S1c |
| | | BR_Vic_0047_2_2011 | PX277616 | DNA-A | ClaI | OxYVV-S1c |
| 104 | 2012 | BR_Vic_0104_4_2012 | PX277617 | DNA-A | EcoRV | OxYVV-S1a |
| 105 | 2012 | BR_Vic_0105_1_2012 | PX277618 | DNA-A | EcoRV | OxYVV-S1a |
| 106 | 2012 | BR_Vic_0106_1_2012 | PX277619 | DNA-A | EcoRV | OxYVV-S1a |
| | | BR_Vic_0106_2_2012 | PX277620 | DNA-A | EcoRV | OxYVV-S1a |
| 107 | 2012 | BR_Vic_0107_1_2012 | PX277621 | DNA-A | EcoRV | OxYVV |
| 111 | 2012 | BR_Vic_0111_1_2012 | PX277622 | DNA-A | EcoRV | OxYVV-S1a |
| | | BR_Vic_0111_2_2012 | PX277623 | DNA-A | EcoRV | OxYVV-S1a |
| 112 | 2012 | BR_Vic_0112_1_2012 | PX277624 | DNA-A | EcoRV | OxYVV-S1c |
| 114 | 2012 | BR_Vic_0114_1_2012 | PX277625 | DNA-A | ClaI | OxYVV-S1b |
| 116 | 2012 | BR_Vic_0116_1_2012 | PX277626 | DNA-A | EcoRV | OxYVV-S1c |

| | | | | | | |
|---|---|---|---|---|---|---|
| 117 | 2012 | BR_Vic_0117_1_2012 | PX277627 | DNA-A | EcoRV | OxYVV-S1c |
| 119 | 2012 | BR_Vic_0119_1_2012 | PX277628 | DNA-A | EcoRV | OxYVV-S1a |
| 120 | 2012 | BR_Vic_0120_1_2012 | PX277629 | DNA-A | EcoRV | OxYVV-S1a |
| 123 | 2012 | BR_Vic_0123_1_2012 | PX277630 | DNA-A | EcoRV | OxYVV-S1a |
| 125 | 2012 | BR_Vic_0125_1_2012 | PX277800 | DNA-A | EcoRV | SiYLCV |
| 126 | 2012 | BR_Vic_0126_1_2012 | PX277631 | DNA-A | EcoRV | OxYVV-S1a |
| | | BR_Vic_0126_2_2012 | PX277763 | DNA-B | EcoRV | OxYVV |
| 128 | 2012 | BR_Vic_0128_1_2012 | PX277632 | DNA-A | EcoRV | OxYVV-S1a |
| 131 | 2012 | BR_Vic_0131_1_2012 | PX277633 | DNA-A | EcoRV | OxYVV-S1a |
| 132 | 2012 | BR_Vic_0132_1_2012 | PX277634 | DNA-A | EcoRV | OxYVV-S1a |
| 133 | 2012 | BR_Vic_0133_1_2012 | PX277635 | DNA-A | ClaI | OxYVV-S1a |
| 136 | 2012 | BR_Vic_0136_1_2012 | PX277636 | DNA-A | ClaI | OxYVV-S1c |
| 201 | 2013 | BR_Vic_0201_1_2013 | PX277637 | DNA-A | ClaI | OxYVV-S1c |
| 206 | 2013 | BR_Vic_0206_1_2013 | PX277638 | DNA-A | PteI | OxYVV-S1a |
| 211 | 2013 | BR_Vic_0211_1_2013 | PX277639 | DNA-A | ClaI | OxYVV-S1a |
| 212 | 2013 | BR_Vic_0212_1_2013 | PX277640 | DNA-A | BssHII | OxYVV-S1a |
| 215 | 2013 | BR_Vic_0215_1_2013 | PX277641 | DNA-A | BssHII | OxYVV-S1a |
| 218 | 2013 | BR_Vic_0218_1_2013 | PX277642 | DNA-A | ClaI | OxYVV-S1a |
| 220 | 2013 | BR_Vic_0220_1_2013 | PX277643 | DNA-A | BssHII | OxYVV-S1a |
| 224 | 2013 | BR_Vic_0224_4_2013 | PX277871 | DNA-B | BssHII | SiYLCV |
| 225 | 2013 | BR_Vic_0225_1_2013 | PX277644 | DNA-A | BssHII | OxYVV-S1a |
| 226 | 2013 | BR_Vic_0226_1_2013 | PX277645 | DNA-A | ClaI | OxYVV-S1a |
| 227 | 2013 | BR_Vic_0227_1_2013 | PX277646 | DNA-A | BssHII | OxYVV-S1a |
| 228 | 2013 | BR_Vic_0228_1_2013 | PX277647 | DNA-A | ClaI | OxYVV-S1a |
| 229 | 2013 | BR_Vic_0229_1_2013 | PX277648 | DNA-A | ClaI | OxYVV-S1a |
| 230 | 2013 | BR_Vic_0230_1_2013 | PX277796 | DNA-B | BssHII | OxYVV |
| 230 | 2013 | BR_Vic_0230_3_2013 | PX277649 | DNA-A | BssHII | OxYVV |
| 230 | 2013 | BR_Vic_0230_4_2013 | PX277650 | DNA-A | BssHII | OxYVV |
| 231 | 2013 | BR_Vic_0231_1_2013 | PX277651 | DNA-A | BssHII | OxYVV-S1c |
| 232 | 2013 | BR_Vic_0232_1_2013 | PX277652 | DNA-A | BssHII | OxYVV-S1a |
| 233 | 2013 | BR_Vic_0233_1_2013 | PX277653 | DNA-A | ClaI | OxYVV-S1a |
| 237 | 2013 | BR_Vic_0237_1_2013 | PX277654 | DAN-A | BssHII | OxYVV-S1a |
| 253 | 2013 | BR_Vic_0253_1_2013 | PX277655 | DNA-A | BssHII | OxYVV-S1a |
| 256 | 2013 | BR_Vic_0256_1_2013 | PX277656 | DNA-A | BssHII | OxYVV-S1a |
| 301 | 2014 | BR_Vic_0301_2_2014 | PX277872 | DNA-B | PteI | SiYLCV |
| 303 | 2014 | BR_Vic_0303_1_2014 | PX277657 | DNA-A | BssHII | OxYVV-S1a |
| | | BR_Vic_0303_2_2014 | PX277658 | DNA-A | BssHII | OxYVV-S1a |
| 304 | 2014 | BR_Vic_0304_06_2014 | PX277873 | DNA-B | ClaI | SiYLCV |

| | | | | | | |
|---|---|---|---|---|---|---|
| 307 | 2014 | BR_Vic_0307_1_2014 | PX277659 | DNA-A | BssHII | OxYVV-S1a |
| 309 | 2014 | BR_Vic_0309_1_2014 | PX277660 | DNA-A | PteI | OxYVV-S1c |
| 310 | 2014 | BR_Vic_0310_1_2014 | PX277661 | DNA-A | ClaI | OxYVV-S1a |
| | | BR_Vic_0310_2_2014 | PX277772 | DNA-B | ClaI | OxYVV-S1a |
| 311 | 2014 | BR_Vic_0311_1_2014 | PX277662 | DNA-A | BssHII | OxYVV-S1a |
| | | BR_Vic_0311_2_2014 | PX277663 | DNA-A | BssHII | OxYVV-S1a |
| 312 | 2014 | BR_Vic_0312_1_2014 | PX277664 | DNA-A | BssHII | OxYVV-S1a |
| 313 | 2014 | BR_Vic_0313_1_2014 | PX277665 | DNA-A | BssHII | OxYVV-S1a |
| 315 | 2014 | BR_Vic_0315_1_2014 | PX277666 | DNA-A | ClaI | OxYVV-S1a |
| 317 | 2014 | BR_Vic_0317_1_2014 | PX277667 | DNA-A | BssHII | OxYVV-S1a |
| 318 | 2014 | BR_Vic_0318_1_2014 | PX277668 | DNA-A | BssHII | OxYVV-S1a |
| 319 | 2014 | BR_Vic_0319_1_2014 | PX277669 | DNA-A | BssHII | OxYVV-S1a |
| 320 | 2014 | BR_Vic_0320_1_2014 | PX277670 | DNA-A | BssHII | OxYVV-S1a |
| 321 | 2014 | BR_Vic_0321_1_2014 | PX277671 | DNA-A | BssHII | OxYVV-S1a |
| 323 | 2014 | BR_Vic_0323_6_2014 | PX277789 | DNA-B | ClaI | OxYVV-S1a |
| | | BR_Vic_0323_1_2014 | PX277672 | DNA-A | ClaI | OxYVV-S1a |
| | | BR_Vic_323_06_2014 | PX277790 | DNA-B | ClaI | OxYVV |
| 325 | 2014 | BR_Vic_0325_1_2014 | PX277673 | DNA-A | PteI | OxYVV-S1a |
| 326 | 2014 | BR_Vic_0326_1_2014 | PX277674 | DNA-A | BssHII | OxYVV-S1a |
| | | BR_Vic_0326_2_2014 | PX277675 | DNA-A | BssHII | OxYVV-S1a |
| 327 | 2014 | BR_Vic_0327_1_2014 | PX277676 | DNA-A | BssHII | OxYVV-S1a |
| 329 | 2014 | BR_Vic_0329_1_2014 | PX277677 | DNA-A | BssHII | OxYVV-S1a |
| 331 | 2014 | BR_Vic_0331_1_2014 | PX277678 | DNA-A | PteI | OxYVV-S1a |
| 334 | 2014 | BR_Vic_0334_1_2014 | PX277679 | DNA-A | EcoRV | OxYVV-S1a |
| 335 | 2014 | BR_Vic_0335_1_2014 | PX277680 | DNA-A | PteI | OxYVV-S1a |
| 337 | 2014 | BR_Vic_0337_1_2014 | PX277681 | DNA-A | PteI | OxYVV-S1a |
| 338 | 2014 | BR_Vic_0338_1_2014 | PX277682 | DNA-A | PteI | OxYVV-S1a |
| 401 | 2016 | BR_Vic_0401_1_2016 | PX277683 | DNA-A | BssHII | OxYVV-S1b |
| 403 | 2016 | BR_Vic_0403_1_2016 | PX277809 | DNA-A | BssHII | SIYLCV |
| | | BR_Vic_0403_2_2016 | PX277810 | DNA-A | BssHII | SIYLCV |
| 404 | 2016 | BR_Vic_0404_3_2016 | PX277875 | DNA-B | BssHII | SiYLCV |
| | | BR_Vic_0404_09_2016 | PX277874 | DNA-B | BssHII | SiYLCV |
| 405 | 2016 | BR_Vic_0405_1_2016 | PX277684 | DNA-A | PteI | OxYVV-S1b |
| | | BR_Vic_0405_2_2016 | PX277685 | DNA-A | PteI | OxYVV-S1b |
| 406 | 2016 | BR_Vic_0406_1_2016 | PX277686 | DNA-A | BssHII | OxYVV-S1b |
| 407 | 2016 | BR_Vic_0407_1_2016 | PX277687 | DNA-A | ClaI | OxYVV-S1b |
| 408 | 2016 | BR_Vic_0408_1_2016 | PX277688 | DNA-A | PteI | OxYVV-S1b |
| 409 | 2016 | BR_Vic_0409_1_2016 | PX277689 | DNA-A | PteI | OxYVV-S1a |

| | | | | | | |
|---|---|---|---|---|---|---|
| | | BR_Vic_0409_2_2016 | PX277690 | DNA-A | PteI | OxYVV-S1a |
| | | BR_Vic_0409_3_2016 | PX277691 | DNA-A | PteI | OxYVV-S1a |
| 410 | 2019 | BR_Vic_0410_1_2016 | PX277692 | DNA-A | PteI | OxYVV-S1a |
| 411 | 2016 | BR_Vic_0411_3_2016 | PX277876 | DNA-B | PteI | SiYLCV |
| 412 | 2016 | BR_Vic_0412_1_2016 | PX277828 | DNA-A | EcoRV | SiYLCV |
| 414 | 2016 | BR_Vic_0414_1_2016 | PX277841 | DNA-A | PteI | SiYLCV |
| | 2016 | BR_Vic_0414_16_2016 | PX277877 | DNA-B | PteI | n.d. |
| 415 | 2016 | BR_Vic_0415_1_2016 | PX277811 | DNA-A | BssHII | SiYLCV |
| | | BR_Vic_415_07_2016 | PX277913 | DNA-B | BssHII | SiYLCV |
| 416 | 2016 | BR_Vic_0416_1_2016 | PX277829 | DNA-A | BssHII | SiYLCV |
| | | BR_Vic_0416_2_2016 | PX277830 | DNA-A | PteI | SiYLCV |
| 418 | 2016 | BR_Vic_0418_2_2016 | PX277878 | DNA-B | BssHII | SiYLCV |
| 419 | 2016 | BR_Vic_0419_07_2016 | PX277879 | DNA-B | PteI | SiYLCV |
| | | BR_Vic_0419_5_2016 | PX277880 | DNA-B | PteI | SiYLCV |
| 421 | 2016 | BR_Vic_0421_1_2016 | PX277842 | DNA-A | EcoRV | SiYLCV |
| 423 | 2016 | BR_Vic_0423_1_2016 | PX277807 | DNA-A | PteI | SiYLCV |
| | | BR_Vic_0423_4_2016 | PX277881 | DNA-B | PteI | SiYLCV |
| 426 | 2016 | BR_Vic_0426_1_2016 | PX277837 | DNA-A | EcoRV | SiYLCV |
| 428 | 2016 | BR_Vic_0428_1_2016 | PX277840 | DNA-A | PteI | SiYLCV |
| | | BR_Vic_0428_2_2016 | PX277832 | DNA-A | PteI | SiYLCV |
| 429 | 2016 | BR_Vic_0429_1_2016 | PX277813 | DNA-A | BssHII | SiYLCV |
| | | BR_Vic_0429_2_2016 | PX277831 | DNA-A | BssHII | SiYLCV |
| 501 | 2017 | BR_Vic_0501_1_2016 | PX277844 | DNA-A | PteI | SiYLCV |
| 502 | 2017 | BR_Vic_0502_1_2017 | PX277812 | DNA-A | PteI | SiYLCV |
| 504 | 2017 | BR_Vic_0504_1_2017 | PX277822 | DNA-A | PteI | SiYLCV |
| 505 | 2017 | BR_Vic_0505_1_2017 | PX277847 | DNA-A | PteI | SiYLCV |
| | | BR_Vic_0505_2_2017 | PX277882 | DNA-B | PteI | SiYLCV |
| | | BR_Vic_0505_7_2017 | PX316469 | DNA-A | PteI | SiYLCV |
| 506 | 2017 | BR_Vic_0506_1_2017 | PX277843 | DNA-A | BssHII | SiYLCV |
| | | BR_Vic_0506_1_2017 | PX277883 | DNA-B | BssHII | SiYLCV |
| 507 | 2017 | BR_Vic_0507_1_2017 | PX277823 | DNA-A | PteI | SiYLCV |
| 508 | 2017 | BR_Vic_0508_1_2017 | PX277693 | DNA-A | EcoRV | OxYVV-S1a |
| 510 | 2017 | BR_Vic_0510_2_2017 | PX277884 | DNA-B | PteI | SiYLCV |
| 511 | 2017 | BR_Vic_0511_1_2017 | PX277814 | DNA-A | PteI | SiYLCV |
| 512 | 2017 | BR_Vic_0512_1_2017 | PX277808 | DNA-A | PteI | SiYLCV |
| | | BR_Vic_0512_5_2017 | PX316470 | DNA-A | PteI | SiYLCV |
| 513 | 2017 | BR_Vic_0513_4_2017 | PX277694 | DNA-A | EcoRV | OxYVV-S1a |
| 514 | 2017 | BR_Vic_0514_1_2017 | PX277695 | DNA-A | PteI | OxYVV-S1b |

| | | | | | | |
|---|---|---|---|---|---|---|
| 515 | 2017 | BR_Vic_0515_1_2017 | PX277696 | DNA-A | PteI | OxYVV-S1a |
| 516 | 2017 | BR_Vic_0516_1_2017 | PX277845 | DNA-A | EcoRV | SiYLCV |
| | | BR_Vic_0516_2_2017 | PX277846 | DNA-A | EcoRV | SiYLCV |
| 517 | 2017 | BR_Vic_0517_1_2017 | PX277697 | DNA-A | PteI | OxYVV-S1a |
| **518** | 2017 | BR_Vic_0518_1_2017 | PX277885 | DNA-A | PteI | OxYVV-S1a |
| | | BR_Vic_0518_1_2017 | PX277698 | DNA-A | PteI | SiYLCV |
| 520 | 2017 | BR_Vic_0520_1_2017 | PX277699 | DNA-A | PteI | OxYVV-S1a |
| 522 | 2017 | BR_Vic_0522_1_2017 | PX277700 | DNA-A | PteI | OxYVV-S1a |
| | | BR_Vic_0522_2_2017 | PX277701 | DNA-A | PteI | OxYVV-S1a |
| 524 | 2017 | BR_Vic_0524_1_2017 | PX277702 | DNA-A | PteI | OxYVV-S1a |
| 525 | 2017 | BR_Vic_0525_1_2017 | PX289945 | DNA-B | PteI | n.d. |
| **527** | 2017 | BR_Vic_0527_1_2017 | PX277824 | DNA-A | PteI | SiYLCV |
| | | BR_Vic_0527_2_2017 | PX277703 | DNA-A | PteI | OxYVV-S1a |
| 529 | 2017 | BR_Vic_0529_1_2017 | PX277704 | DNA-A | PteI | OxYVV-S1a |
| 530 | 2017 | BR_Vic_0530_1_2017 | PX277801 | DNA-A | PteI | SiYLCV |
| 531 | 2017 | BR_Vic_0531_1_2017 | PX277803 | DNA-A | EcoRV | SiYLCV |
| 603 | 2018 | BR_Vic_0603_1_2018 | PX277821 | DNA-A | PteI | SiYLCV |
| | | BR_Vic_0603_5_2018 | PX277886 | DNA-B | PteI | SiYLCV |
| 604 | 2018 | BR_Vic_0604_1_2018 | PX277827 | DNA-A | PteI | SiYLCV |
| 607 | 2018 | BR_Vic_0607_07_2018 | PX246749 | DNA-B | PteI | SimMV |
| 608 | 2018 | BR_Vic_0608_1_2018 | PX277569 | DNA-A | PteI | SimMV-a |
| 610 | 2018 | BR_Vic_0610_1_2018 | PX277839 | DNA-A | PteI | SiYLCV |
| | | BR_Vic_0610_1_2018 | PX277887 | DNA-B | PteI | SiYLCV |
| 612 | 2018 | BR_Vic_0612_07_2018 | PX277888 | DNA-B | PteI | SiYLCV |
| 615 | 2018 | BR_Vic_0615_1_2018 | PX277836 | DNA-A | PteI | SiYLCV |
| 618 | 2018 | BR_Vic_0618_1_2018 | PX277573 | DNA-A | PteI | SimMV-a |
| | | BR_Vic_0618_2_2018 | PX277574 | DNA-A | PteI | SimMV-a |
| 619 | 2018 | BR_Vic_0619_1_2018 | PX277705 | DNA-A | PteI | OxYVV-S1a |
| 620 | 2018 | BR_Vic_0620_1_2018 | PX277706 | DNA-A | PteI | OxYVV-S1a |
| 622 | 2018 | BR_Vic_0622_2_2018 | PX277889 | DNA-B | PteI | SiYLCV |
| | | BR_Vic_0622_1_2018 | PX277802 | DNA-A | PteI | SiYLCV |
| | | BR_Vic_0622_24_2018 | PX277890 | DNA-B | PteI | SiYLCV |
| 623 | 2018 | BR_Vic_0623_1_2018 | PX277707 | DNA-A | PteI | OxYVV-S1a |
| 624 | 2018 | BR_Vic_0624_1_2018 | PX277708 | DNA-A | PteI | OxYVV-S1a |
| 625 | 2018 | BR_Vic_0625_1_2018 | PX277709 | DNA-A | PteI | OxYVV-S1a |
| 626 | 2018 | BR_Vic_0626_1_2018 | PX277849 | DNA-A | PteI | SiYLCV |
| | | BR_Vic_0626_2_2018 | PX277850 | DNA-A | PteI | SiYLCV |
| | | BR_Vic_0626_3_2018 | PX277851 | DNA-A | PteI | SiYLCV |

| | | | | | | |
|---|---|---|---|---|---|---|
| 627 | 2018 | BR_Vic_0627_2018 | PX277580 | DNA-A | PteI | SimMV-b |
| 628 | 2018 | BR_Vic_0628_1_2018 | PX277582 | DNA-A | PteI | SimMV-b |
| 630 | 2018 | BR_Vic_0630_1_2018 | PX277710 | DNA-A | PteI | OxYVV-S1a |
| 631 | 2018 | BR_Vic_0631_1_2018 | PX277711 | DNA-A | PteI | OxYVV-S1a |
| 632 | 2018 | BR_Vic_0632_27_2018 | PX289942 | DNA-B | PteI | SiMV |
| 634 | 2018 | BR_Vic_0634_1_2018 | PX277712 | DNA-A | PteI | OxYVV-S1a |
| 636 | 2018 | BR_Vic_0636_1_2018 | PX277713 | DNA-A | PteI | OxYVV-S1a |
| 637 | 2018 | BR_Vic_0637_1_2018 | PX277805 | DNA-A | PteI | SiYLCV |
| | | BR_Vic_0637_2_2018 | PX277891 | DNA-B | PteI | SiYLCV |
| 639 | 2018 | BR_Vic_0639_1_2018 | PX277577 | DNA-A | PteI | SimMV-a |
| 701 | 2019 | BR_Vic_0701_4_2019 | PX277892 | DNA-B | BssHII | SiYLCV |
| 702 | 2019 | BR_Vic_0702_5_2019 | PX277825 | DNA-A | XhoI | SiYLCV |
| | | BR_Vic_0702_8_2019 | PX277826 | DNA-A | XhoI | SiYLCV |
| 703 | 2019 | BR_Vic_0703_10_2019 | PX277893 | DNA-B | BssHII | SiYLCV |
| 704 | 2019 | BR_Vic_0704_5_2019 | PX277833 | DNA-A | EcoRV | SiYLVC |
| | | BR_Vic_0704_13_2019 | PX277894 | DNA-B | BssHII | SiYLVC |
| 705 | 2019 | BR_Vic_0705_1_2019 | PX277895 | DNA-B | BamHI | SiYLCV |
| | | BR_Vic_0705_2_2019 | PX277896 | DNA-B | BamHI | SiYLCV |
| 706 | 2019 | BR_Vic_0706_4_2019 | PX277834 | DNA-A | PteI | SiYLCV |
| 707 | 2019 | BR_Vic_0707_07_2019 | PX277897 | DNA-B | BssHII | SiYLCV |
| 708 | 2019 | BR_Vic_0708_2_2019 | PX277848 | DNA-B | BssHII | SiYLCV |
| **710** | 2019 | BR_Vic_0710_3_2019 | PX277714 | DNA-A | BssHII | OxYVV-S1a |
| | | BR_Vic_0710_9_2019 | PX277715 | DNA-A | BssHII | OxYVV-S1a |
| | | BR_Vic_0710_10_2019 | PX277575 | DNA-A | BssHII | SimMV-a |
| 712 | 2019 | BR_Vic_0712_1_2019 | PX277576 | DNA-A | BssHII | SimMV-a |
| | | BR_Vic_0712_4_2019 | PX246747 | DNA-B | ClaI | SimMV |
| 713 | 2019 | BR_Vic_0713_9_2019 | PX277852 | DNA-A | BssHII | SiYLCV |
| 714 | 2019 | BR_Vic_0714_1_2019 | PX277854 | DNA-A | BssHII | SiYLCV |
| | | BR_Vic_0714_8_2019 | PX316471 | DNA-A | BssHII | SiYLCV |
| 715 | 2019 | BR_Vic_0715_12_2019 | PX277856 | DNA-A | BssHII | SiYLCV |
| 716 | 2019 | BR_Vic_0716_15_2019 | PX277853 | DNA-A | BssHII | SiYLCV |
| 718 | 2019 | BR_Vic_0718_2_2019 | PX277716 | DNA-A | BssHII | OxYVV-S1a |
| | | BR_Vic_0718_8_2019 | PX277717 | DNA-A | BssHII | OxYVV-S1a |
| 719 | 2019 | BR_Vic_0719_8_2019 | PX277718 | DNA-A | BssHII | OxYVV-S1a |
| | | BR_Vic_0719_2_2019 | PX277794 | DNA-B | BssHII | OxYVV |
| 720 | 2019 | BR_Vic_0720_1_2019 | PX277581 | DNA-A | BssHII | SimMV-b |
| 723 | 2019 | BR_Vic_0723_8_2019 | PX277855 | DNA-A | BssHII | SIYLCV |
| 724 | 2019 | BR_Vic_0724_6_2019 | PX277719 | DNA-A | BssHII | OxYVV-S1a |

| | | | | | | |
|---|---|---|---|---|---|---|
| **725** | 2019 | BR_Vic_0725_6_2019 | PX277720 | DNA-A | BssHII | OxYVV-S1a |
| | | BR_Vic_0725_3_2019 | PX277898 | DNA-B | BssHII | SiYLCV |
| 726 | 2019 | BR_Vic_0726_06_2019 | PX246748 | DNA-B | BssHII | SimM |
| 736 | 2019 | BR_Vic_0736_2_2019 | PX277578 | DNA-A | EcoRV | SimMV-a |
| 741 | 2019 | BR_Vic_0741_4_2019 | PX277721 | DNA-A | BssHII | OxYVV-S1a |
| 742 | 2019 | BR_Vic_0742_4_2019 | PX277857 | DNA-A | BssHII | SiYLCV |
| 801 | 2020 | BR_Vic_0801_1_2020 | PX277722 | DNA-A | ClaI | OxYVV-S1a |
| | | BR_Vic_0801_3_2020 | PX277791 | DNA-B | ClaI | OxYVV |
| 802 | 2020 | BR_Vic_0802_9_2020 | PX277815 | DNA-A | BssHII | SiYLCV |
| **803** | 2020 | BR_Vic_0803_3_2020 | PX277570 | DNA-A | EcoRV | SimMV-a |
| | | BR_Vic_0803_4_2020 | PX277571 | DNA-A | EcoRV | SimMV-a |
| | | BR_Vic_0803_1_2020 | PX277899 | DNA-B | BssHII | SiYLCV |
| | | BR_Vic_0803_5_2020 | PX246744 | DNA-B | BssHII | SimMV |
| | | BR_Vic_0803_2_2020 | PX246745 | DNA-B | EcoRV | SimMV |
| | | BR_Vic_0803_08_2020 | PX246746 | DNA-B | EcoRV | SimMV |
| 804 | 2020 | BR_Vic_0804_1_2020 | PX277817 | DNA-A | BssHII | SiYLCV |
| | | BR_Vic_0804_5_2020 | PX277819 | DNA-A | BssHII | SiYLCV |
| | | BR_Vic_0804_8_2020 | PX277820 | DNA-A | BssHII | SiYLCV |
| | | BR_Vic_0804_2_2020 | PX277900 | DNA-B | ClaI | SiYLCV |
| 805 | 2020 | BR_Vic_0805_3_2020 | PX277792 | DNA-B | ClaI | OxYVV |
| 807 | 2020 | BR_Vic_0807_1_2020 | PX277858 | DNA-A | BssHII | SiYLCV |
| 808 | 2020 | BR_Vic_0808_1_2020 | PX277723 | DNA-A | BssHII | OxYVV-S1b |
| 809 | 2020 | BR_Vic_0809_1_2020 | PX277724 | DNA-A | BssHII | OxYVV-S1e |
| 810 | 2020 | BR_Vic_0810_1_2020 | PX277583 | DNA-A | BssHII | SimMV-b |
| | | BR_Vic_0810_2_2020 | PX277584 | DNA-A | BssHII | SimMV-b |
| 812 | 2020 | BR_Vic_0812_10_2020 | PX277816 | DNA-A | BssHII | SiYLCV |
| | | BR_Vic_0812_2_2020 | PX277901 | DNA-B | ClaI | SiYLCV |
| 814 | 2020 | BR_Vic_0814_1_2020 | PX277585 | DNA-A | BssHII | SimMV-b |
| 815 | 2020 | BR_Vic_0815_06_2020 | PX289943 | DNA-B | ClaI | SiMV |
| 816 | 2020 | BR_Vic_0816_3_2020 | PX277726 | DNA-A | ClaI | OxYVV-S1a |
| | | BR_Vic_0816_1_2020 | PX277725 | DNA-A | EcoRV | OxYVV-S1a |
| | | BR_Vic_0816_3_2020 | PX277765 | DNA-B | EcoRV | OxYVV |
| 818 | 2020 | BR_Vic_0818_1_2020 | PX277838 | DNA-A | EcoRV | SiYLCV |
| | | BR_Vic_0818_8_2020 | PX277818 | DNA-A | EcoRV | SiYLCV |
| | | BR_Vic_0818_3_2020 | PX277835 | DNA-A | BssHII | SiYLCV |
| 819 | 2020 | BR_Vic_0819_9_2020 | PX277859 | DNA-A | BssHII | SiYLCV |
| 831 | 2020 | BR_Vic_0831_1_2020 | PX277902 | DNA-B | ClaI | SiYLCV |
| | | BR_Vic_0831_2_2020 | PX316472 | DNA-A | ClaI | SiYLCV |
| 832 | 2020 | BR_Vic_0832_1_2020 | PX277727 | DNA-A | EcoRV | OxYVV-S1b |

| | | | | | | |
|---|---|---|---|---|---|---|
| | | BR_Vic_0832_2_2020 | PX277728 | DNA-A | EcoRV | OxYVV-S1b |
| | | BR_Vic_0832_4_2020 | PX277729 | DNA-A | ClaI | OxYVV-S1b |
| | | BR_Vic_0832_1_2020 | PX277783 | DNA-B | ClaI | OxYVV |
| | | BR_Vic_0832_2_2020 | PX277784 | DNA-B | ClaI | OxYVV |
| 836 | 2020 | BR_Vic_0836_07_2020 | PX246743 | DNA-B | BssHII | SimMV |
| 837 | 2020 | BR_Vic_0837_5_2020 | PX277572 | DNA-A | EcoRV | SimMV-a |
| | | BR_Vic_0837_1_2020 | PX246742 | DNA-B | BssHII | SimMV |
| | | BR_Vic_0837_13_2020 | PX246741 | DNA-B | BssHII | SimMV |
| 839 | 2020 | BR_Vic_0839_1_2020 | PX277860 | DNA-A | EcoRV | SiYLCV |
| | | BR_Vic_0839_3_2020 | PX277861 | DNA-A | EcoRV | SiYLCV |
| 901 | 2021 | BR_Vic_0901_1_2021 | PX277730 | DNA-A | BssHII | OxYVV-S2 |
| | | BR_Vic_0901_1_2021 | PX277797 | DNA-B | BamHI | OxYVV |
| | | BR_Vic_0901_2_2021 | PX277798 | DNA-B | BamHI | OxYVV |
| **902** | 2021 | BR_Vic_0902_6_2021 | PX277731 | DNA-A | BssHII | OxYVV-S2 |
| | | BR_Vic_0902_10_2021 | PX277903 | DNA-B | BssHII | SiYLCV |
| 903 | 2021 | BR_Vic_0903_1_2021 | PX277732 | DNA-A | BssHII | OxYVV-S1a |
| 904 | 2021 | BR_Vic_0904_10_2021 | PX277733 | DNA-A | ClaI | OxYVV-S1a |
| **905** | 2021 | BR_Vic_0905_1_2021 | PX277734 | DNA-A | EcoRV | OxYVV-S1a |
| | | BR_Vic_0905_2_2021 | PX277904 | DNA-B | ClaI | SiYLCV |
| | | BR_Vic_0905_8_2021 | PX316473 | DNA-A | ClaI | SiYLCV |
| 907 | 2021 | BR_Vic_0907_6_2021 | PX277868 | DNA-A | BssHII | SiYLCV |
| | | BR_Vic_0907_8_2021 | PX277869 | DNA-A | BssHII | SiYLCV |
| 908 | 2021 | BR_Vic_0908_2_2021 | PX277735 | DNA-A | BssHII | OxYVV-S2 |
| 909 | 2021 | BR_Vic_0909_8_2021 | PX277736 | DNA-A | EcoRV | OxYVV-S2 |
| 910 | 2021 | BR_Vic_0910_7_2021 | PX277737 | DNA-A | BssHII | OxYVV-S2 |
| 911 | 2021 | BR_Vic_0911_4_2021 | PX289944 | DNA-A | EcoRV | MaYVV |
| 912 | 2021 | BR_Vic_0912_10_2021 | PX289941 | DNA-A | ClaI | SiMV |
| | | BR_Vic_0912_9_2021 | PX289936 | DNA-B | ClaI | SiMV |
| 913 | 2021 | BR_Vic_0913_4_2021 | PX277579 | DNA-A | ClaI | SimMV |
| 917 | 2021 | BR_Vic_0917_2_2021 | PX277738 | DNA-A | BssHII | OxYVV-S1a |
| | | BR_Vic_0917_1_2021 | PX277799 | DNA-B | ClaI | OxYVV |
| 918 | 2021 | BR_Vic_0918_5_2021 | PX277804 | DNA-A | BssHII | SiYLCV |
| 919 | 2021 | BR_Vic_0919_1_2021 | PX277867 | DNA-A | BssHII | SiYLCV |
| 921 | 2021 | BR_Vic_0921_1_2021 | PX289937 | DNA-A | BssHII | SiMV |
| 923 | 2021 | BR_Vic_0923_8_2021 | PX277862 | DNA-A | BssHII | SiYLCV |
| | | BR_Vic_0923_07_2021 | PX277905 | DNA-B | BssHII | SiYLCV |
| 924 | 2021 | BR_Vic_0924_1_2021 | PX289938 | DNA-A | EcoRV | SiMV |
| | | BR_Vic_0924_6_2021 | PX289939 | DNA-A | EcoRV | SiMV |
| 925 | 2021 | BR_Vic_0925_1_2021 | PX277906 | DNA-B | BssHII | SiYLCV |

| | | | | | | |
|---|---|---|---|---|---|---|
| 926 | 2021 | BR_Vic_0926_6_2021 | PX277806 | DNA-A | BssHII | SiYLCV |
| | | BR_Vic_0926_2_2021 | PX277908 | DNA_B | BssHII | SiYLCV |
| | | BR_Vic_0926_5_2021 | PX277909 | DNA-B | BssHII | SiYLCV |
| | | BR_Vic_0926_10_2021 | PX277907 | DNA-B | BssHII | SiYLCV |
| 928 | 2021 | BR_Vic_0928_2_2021 | PX277910 | DNA-B | BssHII | SiYLCV |
| 1001 | 2022 | BR_Vic_1001_1_2022 | PX277864 | DNA-A | EcoRV | SiYLCV |
| 1002 | 2022 | BR_Vic_1002_2_2022 | PX277911 | DNA-B | EcoRV | SiYLCV |
| 1004 | 2022 | BR_Vic_1004_3_2022 | PX277863 | DNA-A | EcoRV | SiYLCV |
| 1006 | 2022 | BR_Vic_1006_1_2022 | PX277739 | DNA-A | EcoRV | OxYVV-S1a |
| 1008 | 2022 | BR_Vic_1008_3_2022 | PX277870 | DNA-A | EcoRV | SiYLCV |
| **1009** | 2022 | BR_Vic_1009_3_2022 | PX277740 | DNA-A | EcoRV | OxYVV-S1a |
| | | BR_Vic_1009_5_2022 | PX277741 | DNA-A | EcoRV | OxYVV-S1a |
| | | BR_Vic_1009_1_2022 | PX277912 | DNA-B | EcoRV | SiYLCV |
| | | BR_Vic_1009_2_2022 | PX316474 | DNA-A | EcoRV | SiYLCV |
| 1010 | 2022 | BR_Vic_1010_1_2022 | PX277865 | DNA-A | EcoRV | SiYLCV |
| 1012 | 2022 | BR_Vic_1012_1_2022 | PX277742 | DNA-A | EcoRV | OxYVV-S1d |
| | | BR_Vic_1012_2_2022 | PX277764 | DNA-B | EcoRV | OxYVV |
| 1013 | 2022 | BR_Vic_1013_1_2022 | PX277743 | DNA-A | EcoRV | OxYVV-S1d |
| 1014 | 2022 | BR_Vic_1014_5_2022 | PX277744 | DNA-A | EcoRV | OxYVV-S1d |
| | | BR_Vic_1014_06_2022 | PX277767 | DNA-B | EcoRV | OxYVV |
| 1015 | 2022 | BR_Vic_1015_2022 | PX277866 | DNA-A | EcoRV | SiYLCV |
| 1016 | 2022 | BR_Vic_1016_2_2022 | PX277745 | DNA-A | EcoRV | OxYVV-S1d |
| 1018 | 2022 | BR_Vic_1018_8_2022 | PX277746 | DNA-A | EcoRV | OxYVV-S1a |
| 1020 | 2022 | BR_Vic_1020_2_2022 | PX277747 | DNA-A | EcoRV | OxYVV-S1a |
| | | BR_Vic_1020_3_2022 | PX277766 | DNA-B | EcoRV | OxYVV |
| **1021** | 2022 | BR_Vic_1021_8_2022 | PX289940 | DNA-A | EcoRV | SiMV |
| | | BR_Vic_1021_10_2022 | PX277768 | DNA-B | EcoRV | OxYVV |

**Supplementary Table S2.** Number of DNA-A clones of each begomovirus and their respective variants obtained from *Sida acuta* samples collected in Viçosa, MG, from 2011 to 2022.

| **Species-variant** | **2011** | **2012** | **2013** | **2014** | **2016** | **2017** | **2018** | **2019** | **2020** | **2021** | **2022** | **Total** |
|---|---|---|---|---|---|---|---|---|---|---|---|---|
| OxYVV-S1a | 19 | 15 | 17 | 25 | 4 | 11 | 9 | 8 | 3 | 3 | 4 | 118 |
| OxYVV-S1b | 2 | 1 | 0 | 0 | 6 | 1 | 0 | 0 | 4 | 0 | 0 | 14 |
| OxYVV-S1c | 10 | 4 | 3 | 1 | 0 | 0 | 0 | 0 | 0 | 0 | 0 | 18 |
| OxYVV-S1d | 0 | 0 | 0 | 0 | 0 | 0 | 0 | 0 | 0 | 0 | 4 | 4 |
| OxYVV-S1e | 0 | 0 | 0 | 0 | 0 | 0 | 0 | 0 | 1 | 0 | 0 | 1 |
| OxYVV-S2 | 0 | 0 | 0 | 0 | 0 | 0 | 0 | 0 | 0 | 6 | 1 | 7 |
| SiYLCV | 0 | 1 | 0 | 0 | 14 | 15 | 9 | 12 | 13 | 7 | 6 | 77 |
| SiMMV-a | 0 | 0 | 0 | 0 | 0 | 0 | 4 | 3 | 3 | 1 | 0 | 11 |
| SiMMV-b | 0 | 0 | 0 | 0 | 0 | 0 | 2 | 1 | 3 | 0 | 0 | 6 |
| SiMV | 0 | 0 | 0 | 0 | 0 | 0 | 0 | 0 | 0 | 4 | 1 | 5 |
| MaYSV | 0 | 0 | 0 | 0 | 0 | 0 | 0 | 0 | 0 | 1 | 0 | 1 |
| Total | 31 | 21 | 20 | 26 | 24 | 27 | 24 | 24 | 26 | 22 | 16 | 262 |

**Supplementary Table S3.** Proportion of non-synonymous to synonymous substitutions for DNA-A genes of the begomoviruses infecting *Sida acuta* in Viçosa, MG, and sites under positive or negative selection according to three maximum likelihood-based methods.

| **Gene** | **Tajima's D** | **Fu & Li's** | | **dN/dS** | **SLAC** | | **MEME** | | **FUBAR** | |
|---|---|---|---|---|---|---|---|---|---|---|
| | | ***F**** | ***D**** | | **Positive** | **Negative** | **Positive** | **Negative** | **Positive** | **Negative** |
| **OxYVV** | | | | | | | | | | |
| *CP* | -1,351 | -1.609 | -1.387 | 0.129 | - | 18, 25, 34, 37, 71, 72 86, 94, 137, 149 154, 156, 177, 209, 217, 225 245, 249 | 31 | - | - | 18, 20, 25, 32, 34, 37, 47, 49, 50, 53, 71, 72, 73, 83, 86, 94, 113, 137, 149, 154, 156, 158, 173, 177, 182, 191, 209, 217, 218, 223, 224, 225, 237, 245, 249 |
| *Rep* | -1.356 | -2,45849* | -2,673* | 0.259 | | 40, 70, 72, 78, 116, 144, 151, 157, 186, 187, 220, 251, 274, 321, 324 | 60, 98, 123, 214 | | - | 17, 39, 40, 62, 70, 72, 78, 110, 115, 116, 126, 143, 144, 151, 157, 166, 178, 184, 186, 187, 194, 220, 223, 235, 251, 260, 274, 314, 321, 324, 332, 347 |
| *TrAP* | -1.396 | -2.904* | -3.201* | 0.534 | - | 2, 15, 121 | 92, 94 | - | 92, 119 | 2, 15, 33, 64, 121 |
| *Ren* | -.1264 | -2.216 | -2.324 | 0.379 | | 35, 47, 73, 74, 103 | 126 | | | 27, 31, 35, 42, 47, 73, 74 102, 103, 122, |
| *AC4* | -1.7233 | -1.048 | -1.611 | 1.35 | - | 58 | 6, 19 | - | 9, 17, 75 | 58, 81 |
| *MP* | 1.1025 | 0,8427 | 0,4650 | 0.113 | | 20, 159, 182 | 233 | | - | 159, 182, 20, 179, 71, 215, 123, 66, 120, 124, 220, 62, 129, 260, 10, 151, 35, 14, 63, 84, 85, 112, 161, 203, 97, 171, 206, 278, 291, 277, 140, 2, 180, 82, 148, 139, 193 |
| *NSP* | 0,5616 | 0,0510 | -0,2724 | 0.275 | - | 48, 115 | 120, 160 | | 120 | 48, 63, 95, 110, 115, 129, 133, 145, 215 |
| **SiYLCV** | | | | | | | | | | |

| | | | | | | | | | | |
|---|---|---|---|---|---|---|---|---|---|---|
| *CP* | 0,3621 | -0,56344 | -1,0069 | 0.151 | | 80, 94, 155,136 | 81 | | 10 | 49, 80, 83, 90, 94, 127, 136, 155, 182, 190, 222, 239 |
| *Rep* | -1,1275 | -1,6539 | -1,5428 | 0.468 | | 52, 125 | 172 | | 44, 172, 181 | 12, 23, 43, 52, 53, 57, 125, 187, 259, 332 |
| *TrAP* | -1,9271* | -2,4176 | -2,0817 | 0.573 | - | - | - | - | 14, 129 | 34, 42, 99, 103 |
| *Ren* | -1,6125* | -1,3532 | -0,8311 | 0.319 | - | - | - | - | - | 122, 130 |
| *AC4* | -1,8756* | -1,7871 | -1,2895 | 1.13 | - | - | - | - | - | - |
| *MP* | 0,5227 | 0,8965 | 0,9076 | 0.154 | - | 41, 43, 66, 149, 162 | 32 | - | | 7, 14, 20, 41, 43, 58, 66, 75, 82, 94, 113, 126, 127, 137, 149, 162, 170, 172, 184, 202, 211, 217 |
| *NSP* | -1,0070 | -0,7648 | -0,4408 | 0.319 | - | 201 | - | - | - | 160, 174, 186, 199, 201, 211 |

**Supplementary figure legends**

**Suppl. Figure S1.** *Sida acuta* plants growing in a suburban area near Viçosa, MG, Brazil, displaying symptoms of begomovirus infection.

**Suppl. Figure S2.** Aspects of the area near Viçosa, MG, Brazil, where *Sida acuta* samples were collected from 2011 to 2022. **A, B.** Satellite images of the area and the surrounding region. The red circles indicate the area were samples were collected. **C, D.** General aspect of the area were samples were collected. Images obtained in 2024.

**Suppl. Figure S3.** Pairwise nucleotide sequence identity matrix of Oxalys yellow vein virus (OxYVV) isolates obtained in this work. The colored bar at the left indicates the two strains and five variants identified based on the cut-off values determined by the *Geminiviridae* Study Group of the International Committee on Taxonomy of Viruses (ICTV) (Brown et al., 2015).

**Suppl. Figure S4.** Pairwise nucleotide sequence identity matrix of Sida yellow leaf curl virus (SiYLCV) isolates obtained in this work.

**Suppl. Figure S5.** Pairwise nucleotide sequence identity matrix of Sida micrantha mosaic virus (SimMV) isolates obtained in this work. The colored bar at the left indicates the two strains identified based on the cut-off values determined by the *Geminiviridae* Study Group of the International Committee on Taxonomy of Viruses (ICTV) (Brown et al., 2015).

**Suppl. Figure S6.** Diversity accumulation curves for different sample sizes. The shaded background represents 95% confidence intervals of Hill numbers calculated for the three diversity orders (0, 1 and 2) estimated for each year of collection.

**Suppl. Figure S7.** Evolutionary and demographic dynamics of the Oxalis yellow vein virus (OxYVV) and Sida yellow leaf curl virus (SiYLCV) coat protein (*CP*) genes. **A.** Maximum clade credibility (MCC) phylogenetic tree for the OxYVV *CP* gene. The numbers on the branches indicate the time to the most recent common ancestor. The identified OxYVV variants (S1a-S1e and S2) are shown in distinct colors in the side bar. **B.** MCC tree for the SiYLCV *CP* gene. **C.** Demographic history under a coalescent framework (effective population size, Ne) inferred using the Bayesian Skygrid model for the OxYVV and SiYLCV *CP* genes. The lines represent the median estimates, and the shaded areas correspond to the 95% highest posterior density (HPD) intervals.

**Suppl. Figure S8.** Evolutionary and demographic dynamics of the Oxalis yellow vein virus (OxYVV) and Sida yellow leaf curl virus (SiYLCV) replication-associated protein (*Rep*) genes. **A.** Maximum clade credibility (MCC) phylogenetic tree for the OxYVV *Rep* gene. The numbers on the branches indicate the time to the most recent common ancestor. The identified OxYVV variants (S1a–S1e and S2) are shown in distinct colors in the side bar. **B.** MCC tree for the SiYLCV *Rep* gene. **C.** Demographic history under a coalescent framework (effective population size, Ne) inferred using the Bayesian Skygrid model for OxYVV and SiYLCV *Rep* genes. The lines represent the median estimates, and the shaded areas correspond to the 95% highest posterior density (HPD) intervals.

**Suppl. Figure S1**

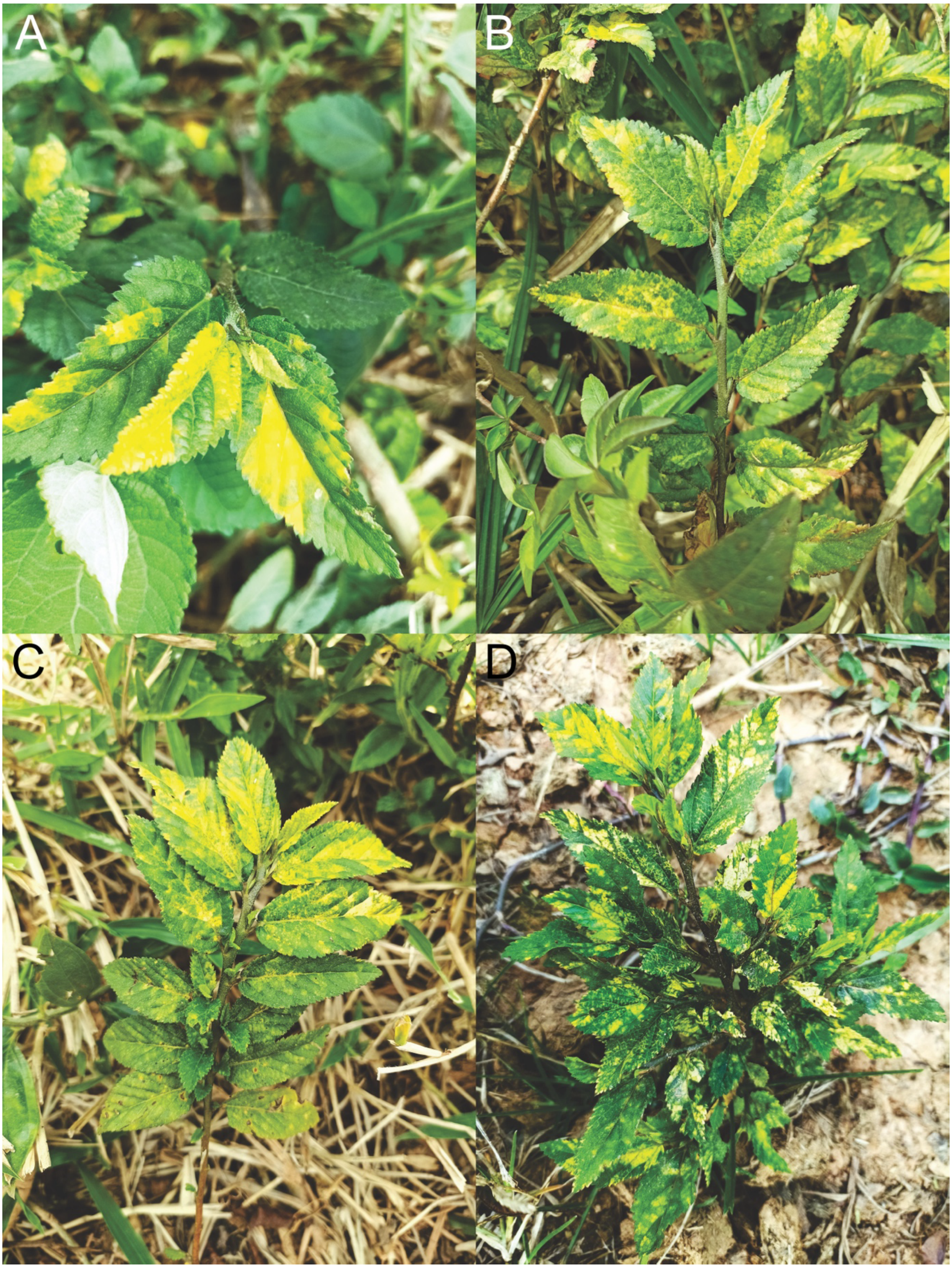

**Suppl. Figure S2**

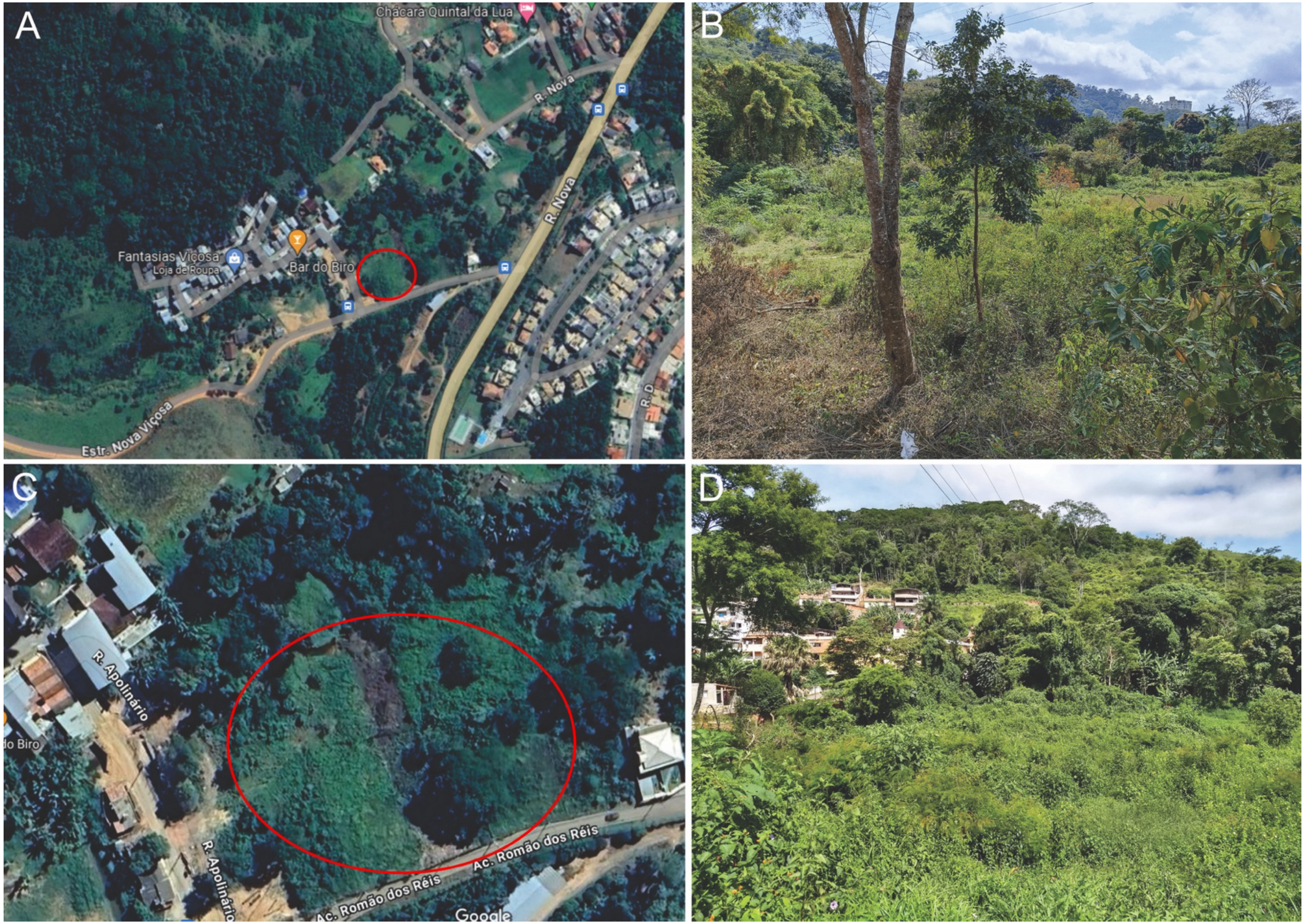

### Suppl. Figure S3

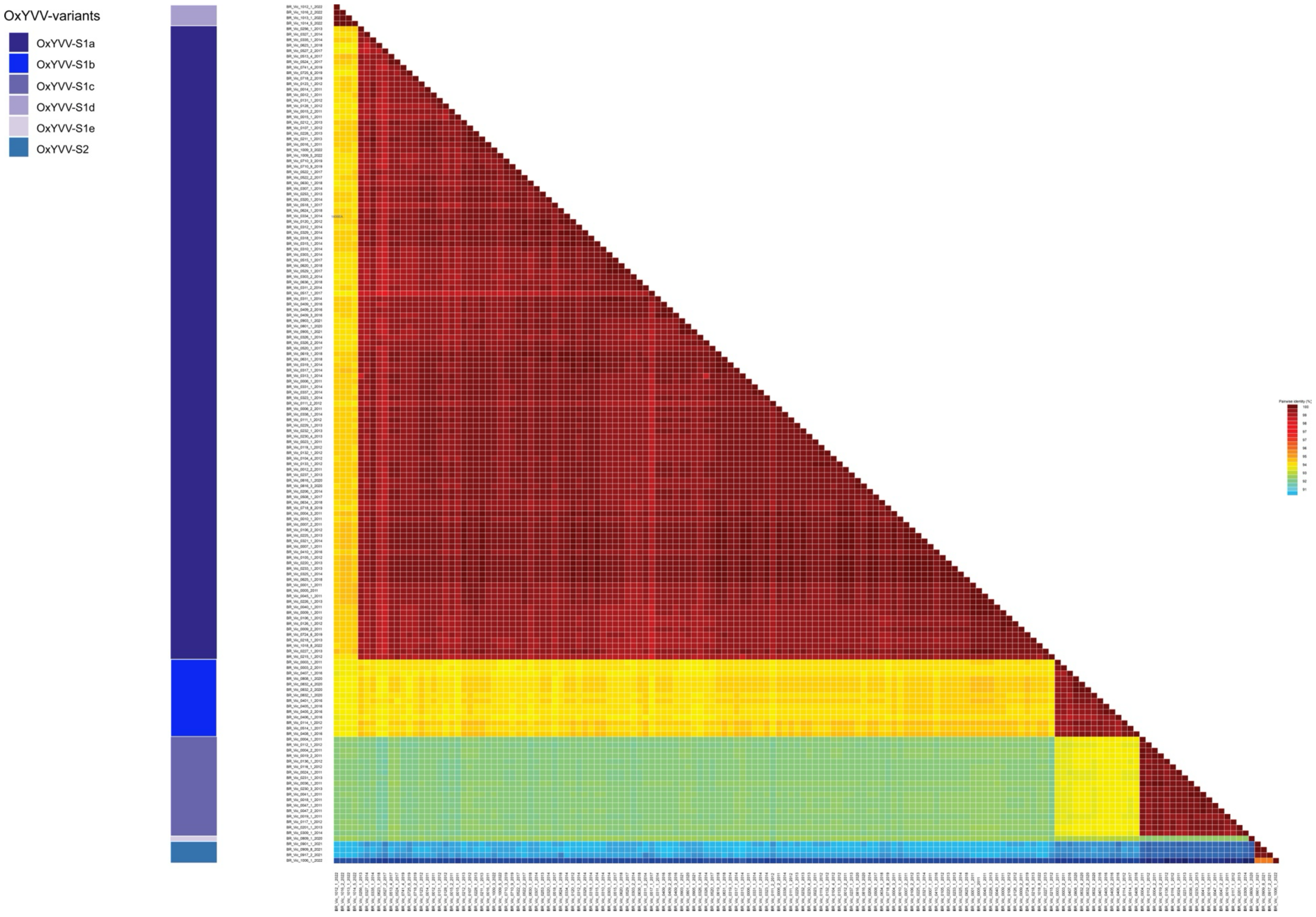

**Suppl. Figure S4**

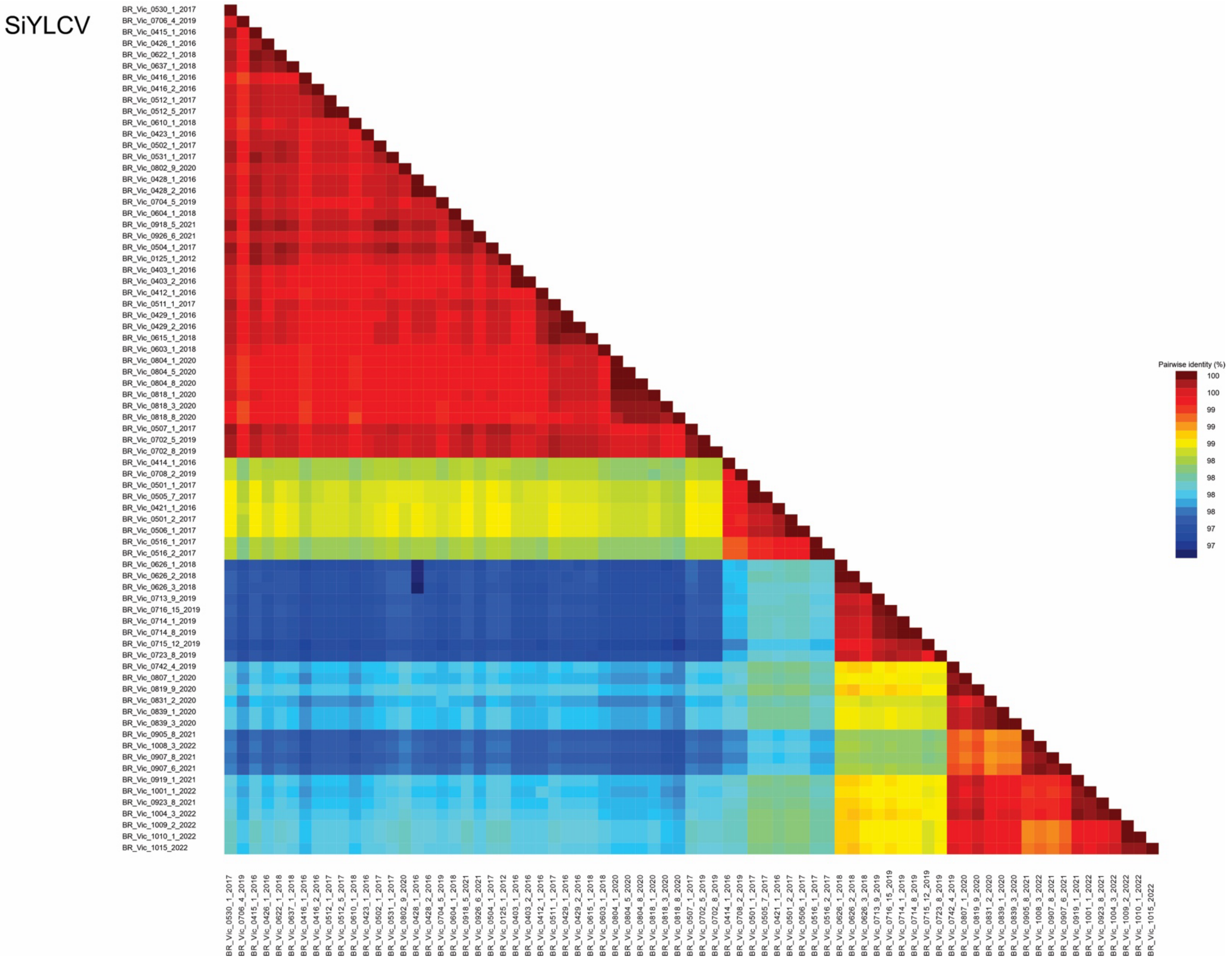

**Suppl. Figure S5**

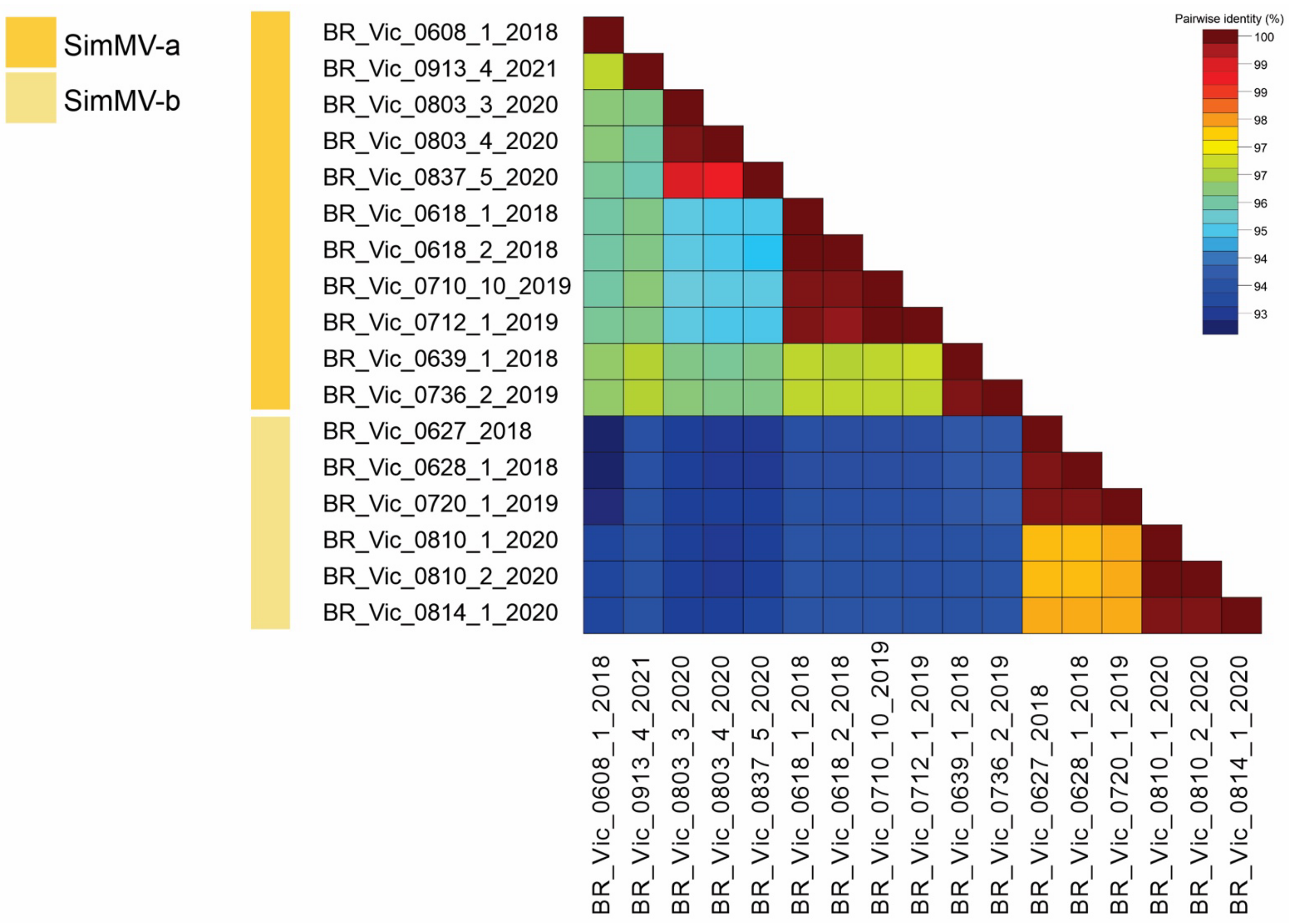

**Suppl. Figure S6**

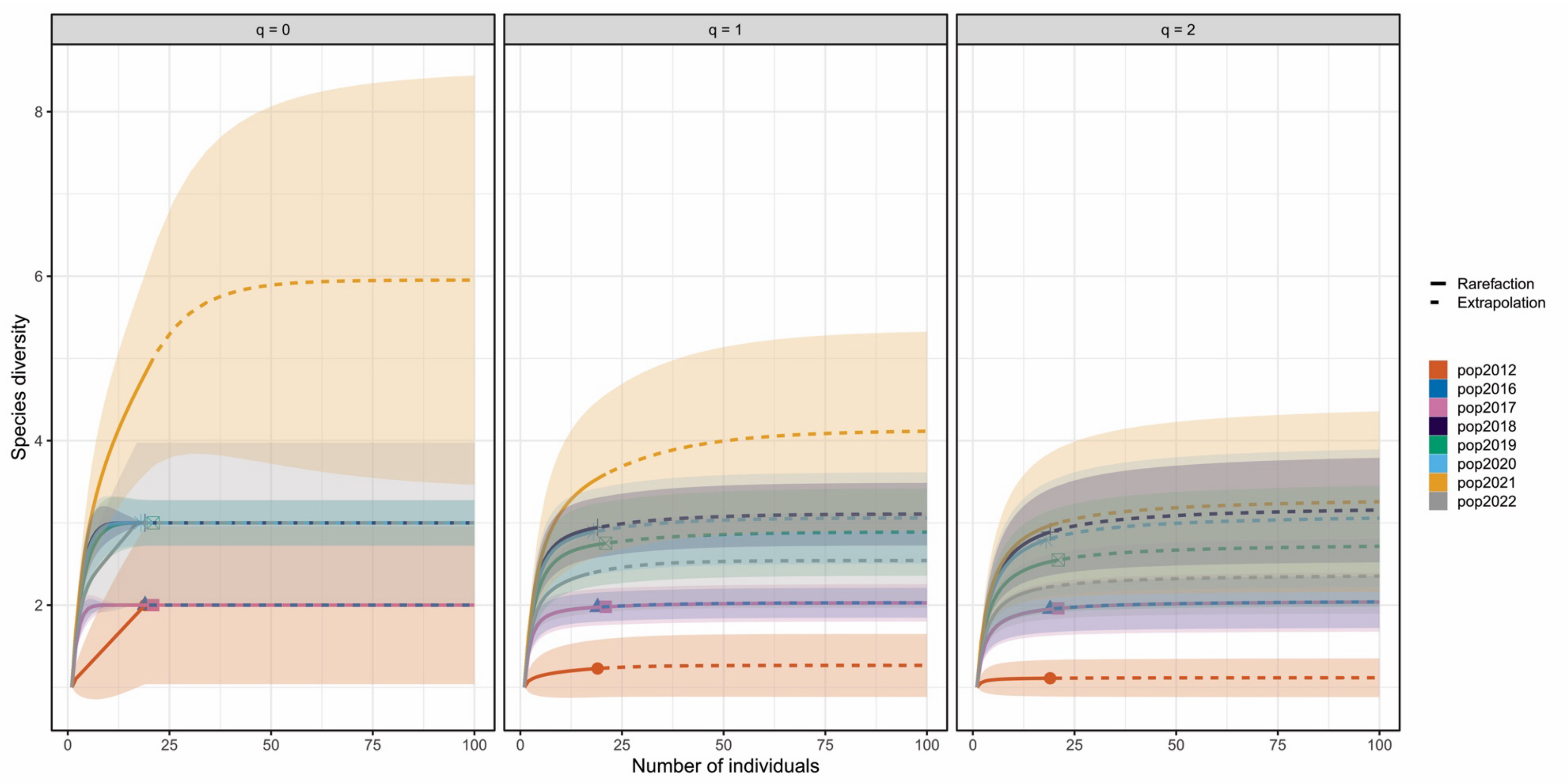

**Suppl. Figure S7**

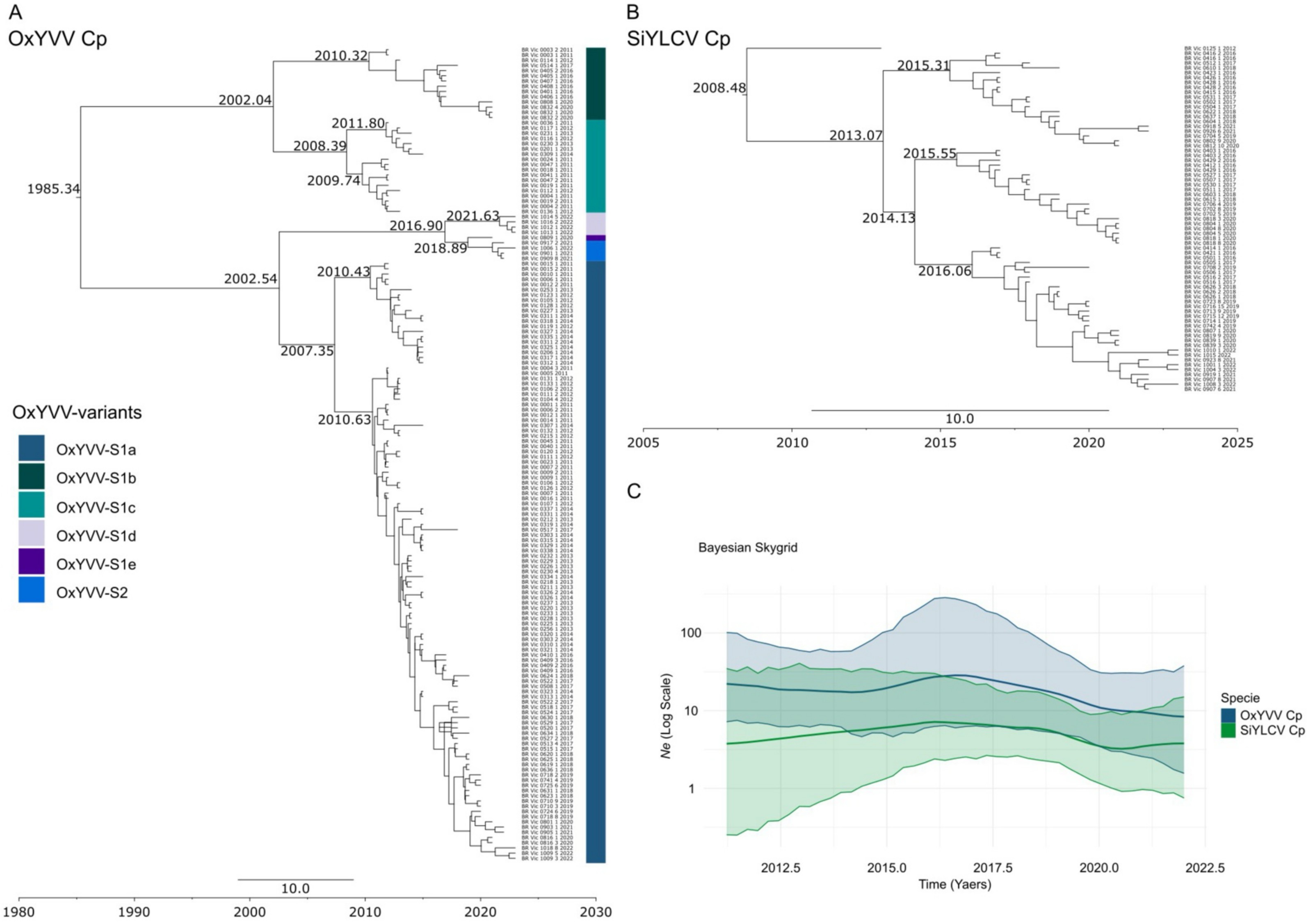

### Suppl. Figure S8

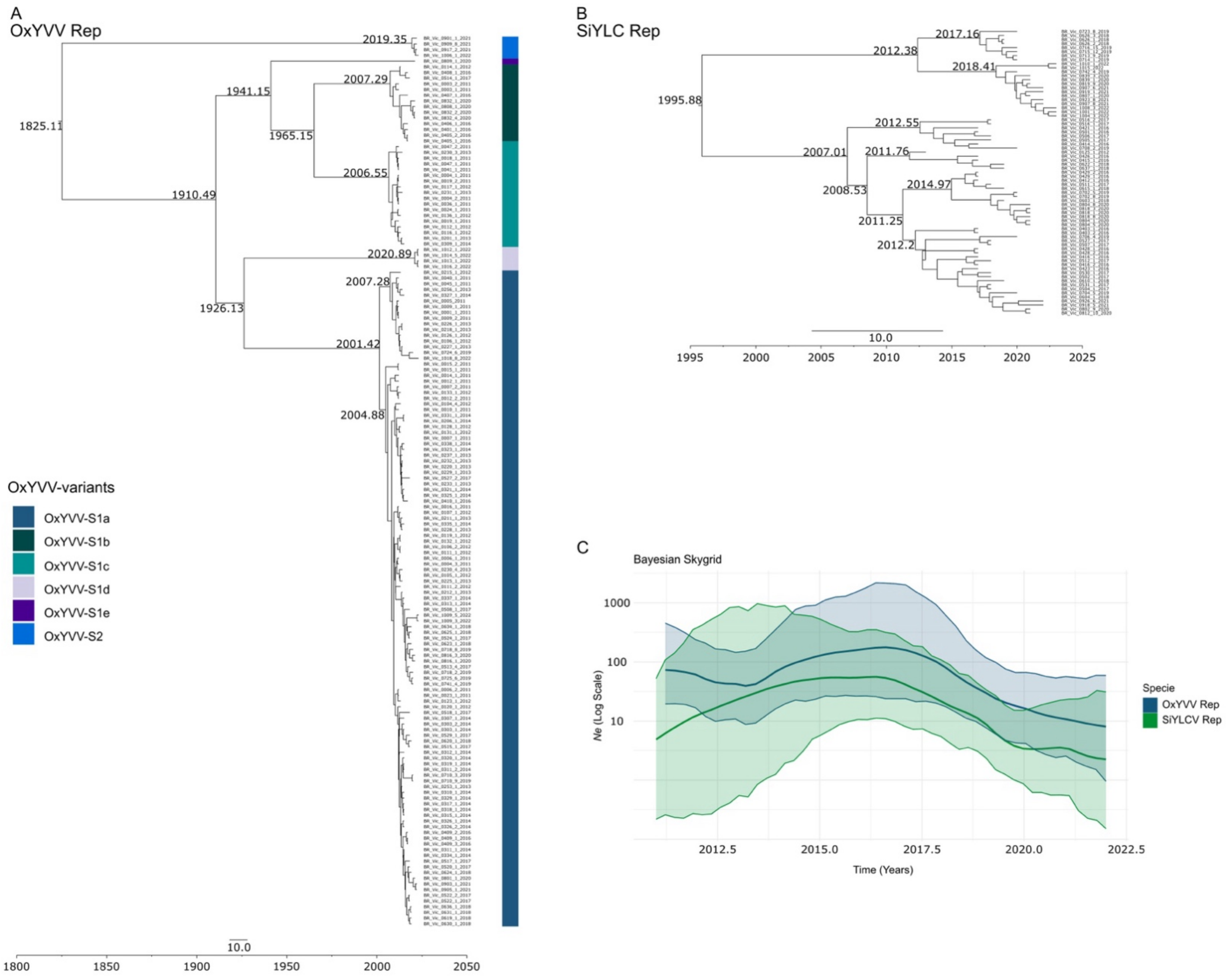
